# Social Media and citizen science provide valuable data for behavioural ecology research: Are cuttlefish using pursuit-deterrent signals during hunting?

**DOI:** 10.1101/760926

**Authors:** Dražen Gordon, Philip Pugh, Gavan M Cooke

## Abstract

Obtaining robust, analysable data sets from wild marine animals is fraught with difficulties, dangers, expense, often without success. Scientists are becoming increasingly reliant on citizen scientists to help fill in gaps where they exist, especially in the area of biodiversity. Here, uniquely, we use social media and citizen science videos to investigate the behavioural ecology of hunting in five cuttlefish species – *Metasepia pfefferi* (N = 24), *Sepia apama* (N = 13), *Sepia latimanus* (N = 8), *Sepia officinalis* (N = 17), and *Sepia pharaonis* (N = 23). We find that hunting strategies and prey type differ between species as do the types of behaviours used by the five species studied here. We also use kinematic permutation analysis to elucidate chains of behaviours, finding that cuttlefish significantly use a mixture of predator behaviours but also prey-like behaviours, such as warning signals and possibly even a ‘pursuit-deterrent signal’ during the final moments of hunting. We also show and discuss significant intraspecific differences.

## Introduction

All animals share the obligation for nourishment as it enables them to grow, survive and reproduce^1^. Animals can be either ‘predators’ or ‘prey’, though many are both^2^. Predators and prey are in an evolutionary arms race, where species-specific fitness enhancing traits are attained via natural selection, since not being eaten or managing to eat is clearly highly adaptive^3^.

Optimal Foraging Theory^4,5^ states that predators should strive to maximise net energy intake and individual fitness through profitable prey selection, and thus predators have evolved numerous different hunting strategies^5,6^. In turn, prey are under strong selection pressure to avoid being eaten and many species have evolved “pursuit-deterrent signals”^7^, that can either express an individual’s fitness, and thus unprofitability as a target of predation (e.g. high likelihood of evading predation), or are seemingly intimidating/startling, distracting, or dissuading for predators^7–9^. For example, some terrestrial arthropods stridulate specialised organs or secrete noxious/distasteful substances, whereas social ungulates (Bovidae, Cervidae, and Antilocapridae) perform stotting displays when a predator stalks them^7,9,10^. Like primary consumers, mesopredators are generally more susceptible to predation during periods of foraging and food handling, as they devote less time to attentive environmental scanning^7,11^. The timing of pursuit-deterrent signal exhibition is thus vital to a target’s fitness and chance of survival^7^. Discouraging predators prior to pursuit or attack helps to avoid potentially high energy expenditure and physical damage during fight or flight situations and/or the loss of food whilst avoiding being eaten themselves. To our knowledge, all presently documented cases of predator-prey interactions describing pursuit-deterrent signals have focused on the prey species, rather than a predator that has lowered its guard to focus on the act of hunting^7–11^.

Almost all cephalopods (Mollusca: cuttlefish, squid, octopus and nautilus) are voracious meso-predators that hunt a diversity of prey – primarily crustaceans, teleost fish, and indeed other cephalopod species^12,13^. An advanced non-centralised nervous system, coupled to neuromuscular and hydrostatic dermal units requires the neural complexity that appears to also provide cephalopods high levels of cognition, significant (including episodic) memory, complex sensory systems (chemo-, mechano-, and photo-receptors), and real-time body pattern manipulation (i.e. deimatic displays). All these attributes are used to locate, identify, and engage prey^13^. Additionally, these unique adaptations are used to determine, and avoid or deter seal, dolphin, shark, and bony fish predators^14,15^ via camouflage^16^ or warning displays^15^.

Hunting mode repertoires vary within the Sepiidae, mature (non-toxic) *Sepia spp*. (e.g., *Sepia apama, Sepia officinalis*, and S*epia pharaonis*) are thought to hunt within short time-periods, incorporating inconspicuous body patterns (i.e. disruptive, mottled, and uniformly stippled) to minimise risk of predation^14,20–23^. *Sepia latimanus* adopts a different strategy, directing conspicuous ‘dynamic passing wave’ patterns towards prey^20^. *Metasepia* spp. (>2 species) forage by ‘crawling’ along the substratum whilst exhibiting aposematic (e.g. yellow or red) colours at all times^24–26^ – signalling their toxicity^23,27^. Hunting strategies of the six *Sepiella spp*., are unstudied^28–30^.

Studying marine animal behaviour is difficult, expensive, often dangerous and uncertain of success, taking many years to gather significant data sets. To counter this, we employed the public to obtain robust data. ‘Citizen science’ broadly refers to public participation and engagement in scientific projects^31^, though biological citizen science has almost exclusively focused on measuring biodiversity^20,31^. In addition to proactive or retroactive data requests, there are numerous animal observations available via Facebook, Twitter, Instagram, YouTube and other social media which we utilised to provide useful data to fill knowledge-gaps in behavioural ecology processes. Cuttlefish, and cephalopods more generally, are a case in point. Field observations are, compared to laboratory studies, relatively rare for cephalopods^16,32,33^ and so provide an excellent model for using unusual data sources.

This study tests: (1) the hypothesis that retroactively gathered citizen science and unsolicited social media observational data can be used to elucidate the behavioural ecology of hunting in five Sepiid species (species: *M. pfefferi, S. apama, S. latimanus, S. officinalis*, and *S. pharaonis*); and (2) the hypothesis that cephalopods use species specific hunting behaviours.

## Materials and methods

### Data acquisition and variables

Videos containing wild feeding events were acquired from “The Cephalopod Citizen Science Project”^34^ and online media sharing websites: Arkive^35^, Shutterstock^36^, NatureFootage^37^, NaturePictureLibrary^38^, Vimeo^39^, Footage.net^40^, and YouTube^41^. Pre-recorded feeding events were filmed by members of the public over the last 10 years covering a large geographic area (SW England to SE Australia – see Figure 1) and varied in the number of observations and environmental hunting conditions. Variables included: subject maturity (all adult); subject sex (unidentified); time of feeding event (night or day); target prey item species; hunting strategy (foraging, ambush, etc.); habitat; and success of prey seizure. We also noticed a wide range of signals deployed by cuttlefish during hunting. Videos were only included if they contained a complete sequence of hunting, from prey detection to prey attack. Length of videos varied (see Table 2) as did average numbers of behaviours per attack sequence, which represents real world circumstances. Comparisons for length of foraging time, length of attack sequences or length of specific behaviours, and other basic inferential statistics, were not attempted due to the difficulty of controlling for video sequence length. Instead, we chose to employ novel approaches to elucidate differences and relationships between five cuttlefish species during hunting behaviour. Whilst citizen science sourced media enabled quick and accessible data collection, it also had limitations. We did not compare day and night (i.e. dark) feeding events, both events took place under illuminated conditions (i.e. SCUBA divers use bright torches at night). The diver’s presence had an unknown influence on cuttlefish predation success and feeding behaviour. For example, warning displays, including possible pursuit-deterrent signals, may have been directed towards divers, although videos with displays obviously directed at the divers were omitted. Predator presence in peripheral vicinities of footage was unknown. However, all these issues could be said of scientists collecting data themselves and is not unique to the untrained public. The species studied here had limited videos (and highly variable observation times – see Table 2) to choose from, and we excluded many because they were incomplete or had uncertain cuttlefish species within them. Lastly, editorial influences and poor video quality inhibited prey species identification and incurred missed behavioural recording, forcing us to reject them from our analysis.

**Table 1.**
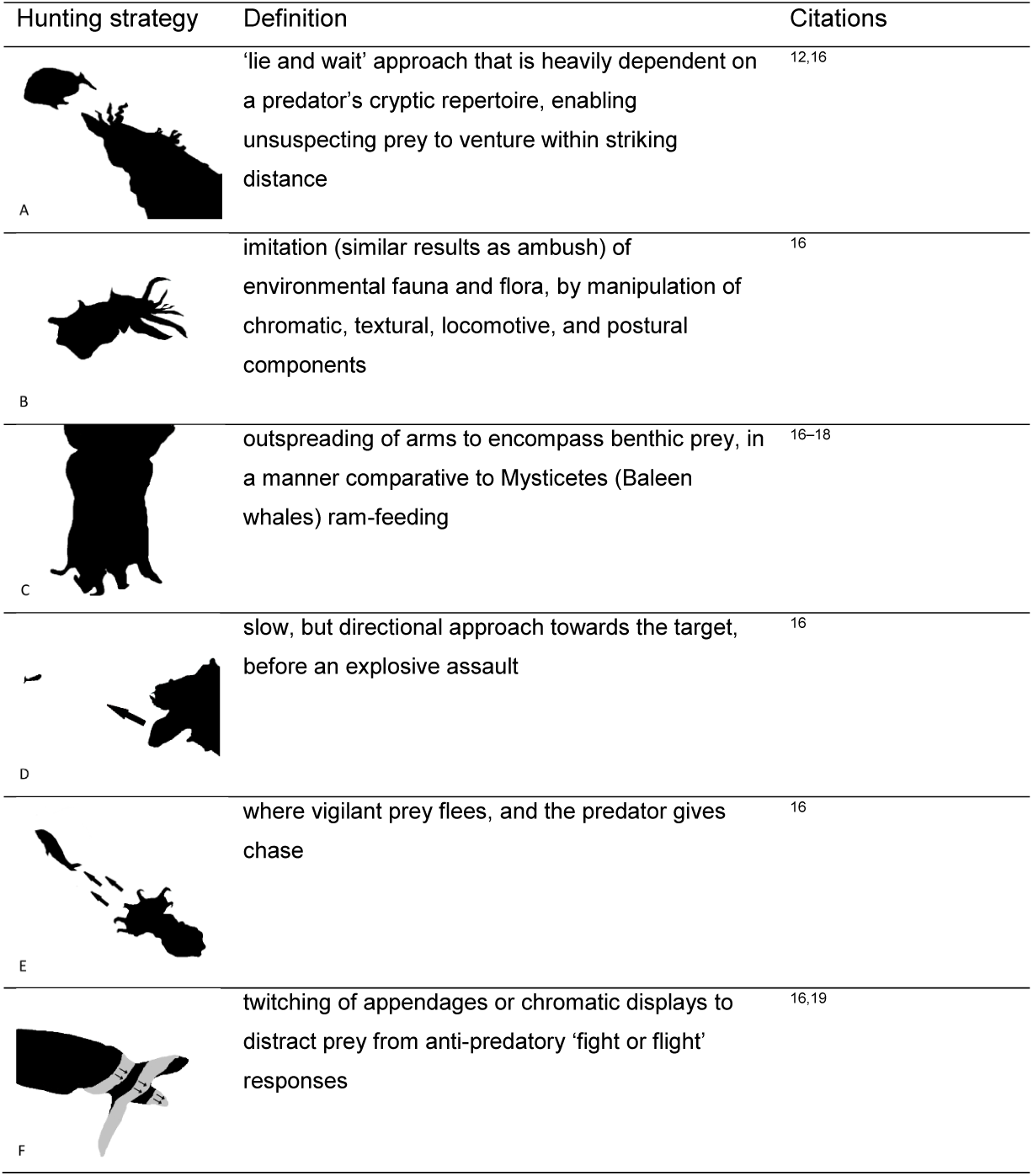
Hunting strategies known to be used by cuttlefish: ambush (A), mimicry (B), speculative hunting (C), stalking (D), guided-pursuit (E), and luring or directive mark (example used: “rhythmic passing waves”) (F).

| Hunting strategy | Definition | Citations |
| --- | --- | --- |
| A | 'lie and wait' approach that is heavily dependent on a predator's cryptic repertoire, enabling unsuspecting prey to venture within striking distance | 12,16 |
| B | imitation (similar results as ambush) of environmental fauna and flora, by manipulation of chromatic, textural, locomotive, and postural components | 16 |
| C | outspreading of arms to encompass benthic prey, in a manner comparative to Mysticetes (Baleen whales) ram-feeding | 16–18 |
| D | slow, but directional approach towards the target, before an explosive assault | 16 |
| E | where vigilant prey flees, and the predator gives chase | 16 |
| F | twitching of appendages or chromatic displays to distract prey from anti-predatory 'fight or flight' responses | 16,19 |

**Table 2.** Number of observations, total behaviours, duration of hunting events and number of ‘flash upon predation’, per species

|  | *M. pfefferi* | *S. apama* | *S. latimanus* | *S. officinalis* | *S. pharaonis* |
| --- | --- | --- | --- | --- | --- |
| No. behaviours recorded | 80 | 57 | 72 | 83 | 74 |
| No. observations | 24 | 13 | 8 | 17 | 23 |
| Avg. Obs. Duration (seconds) | 21.419 ± 4.199 | 6.014 ± 1.415 | 27.214 ± 6.285 | 17.512 ± 3.625 | 8.426 ± 2.370 |
| No. Flash<br>upon<br>predation | 20 | 5 | 6 | 11 | 21 |

**Table 3.** Prey taxa: lifestyle (benthic or pelagic) percentages and observed averages (a); predator evasion percentages and experimental averages (b); and homogeneity of predated taxa by species (one-way classification chi-square). Unidentifiable prey items were excluded from analysis: *M. pfefferi* (n = 3), *S. apama* (n = 1), *S. latimanus* (n = 2), *S. officinalis* (n = 2), and *S. pharaonis* (n = 3) (c).

| (a) | N | Predated Osteichthyes lifestyle |  | Predated Crustacea lifestyle % |  |
| --- | --- | --- | --- | --- | --- |
|  |  | % |  |  |  |
|  |  | Benthic | Pelagic | Benthic | Pelagic |
| *M. pfefferi* | 21 | 84.6 | 15.4 | 100 | 0 |
| *S. apama* | 12 | 25 | 75 | 0 | 0 |
| *S. latimanus* | 8 | 50 | 50 | 100 | 0 |
| *S. officinalis* | 15 | 60 | 40 | 100 | 0 |
| *S. pharaonis* | 18 | 13.3 | 86.7 | 100 | 0 |
| Avg. % |  | 46.6 ± 12.7 | 53.4 ± 12.7 | 100 ± 0 | 0 ± 0 |

| (b) | N | Prey item % |  | Successful prey seizure % |  |
| --- | --- | --- | --- | --- | --- |
|  |  | Osteichthyes | Crustacea | Osteichthyes | Crustacea |
| *M. pfefferi* | 21 | 61.9 | 38.1 | 69.2 | 100 |
| *S. apama* | 12 | 100 | 0 | 75 | 0 |
| *S. latimanus* | 8 | 75 | 25 | 50 | 100 |
| *S. officinalis* | 15 | 66.7 | 33.3 | 70 | 60 |
| *S. pharaonis* | 18 | 83.3 | 16.7 | 60 | 100 |
| Avg. % |  | 77.4 ± 6.7 | 22.6 ± 6.7 | 64.9 ± 4.4 | 90 ± 10 |

| (c) | Homogeneity of dietary composition |  |  |
| --- | --- | --- | --- |
|  | N | X<sup>2</sup> | p |
| *M. pfefferi* | 21 | 1.19 | 0.275 |
| *S. apama* | 12 | 12 | <0.001 |
| *S. latimanus* | 8 | 2 | 0.157 |
| *S. officinalis* | 15 | 1.67 | 0.197 |
| *S. pharaonis* | 18 | 8 | 0.005 |

**Figure 1.**
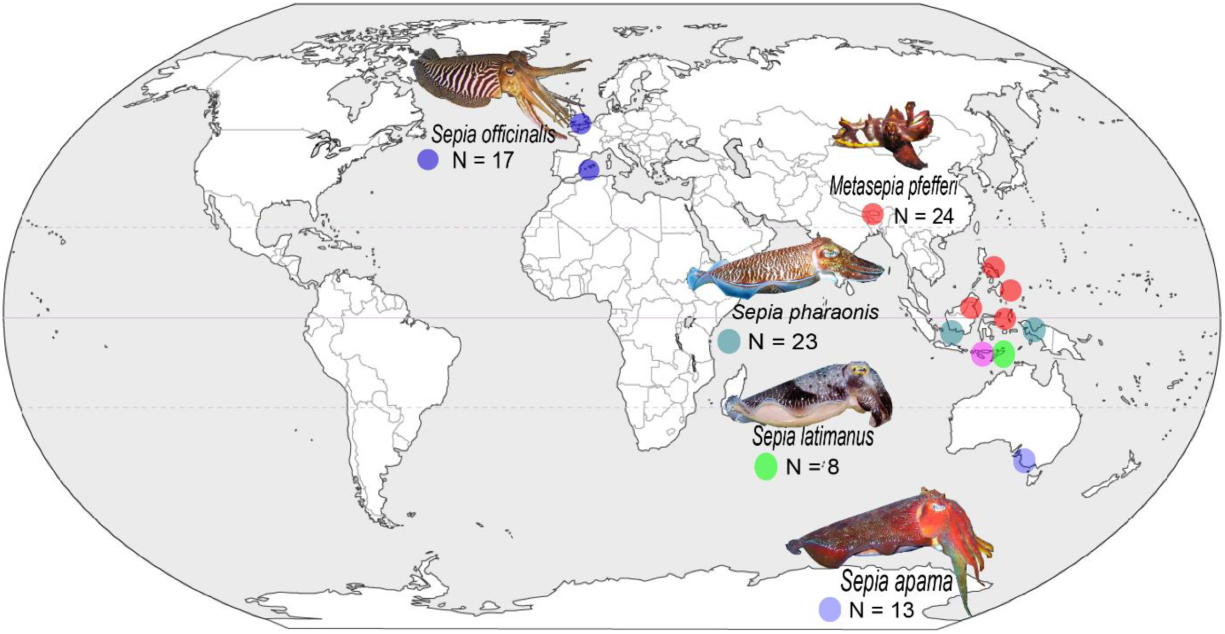
General geographic locations of citizen science and social media sources, for the five cuttlefish species studied here, showing solitary cuttlefish feeding events. N = number of videos per species that were analysed.

### Ethogram creation

Species specific fine scale ethograms were created to catalogue behaviours expressed amongst species where observations exist in enough numbers for analysis. Ethograms were designed to be observation-specific, and therefore excluded pre-documented/unseen behaviours (see Supplementary Table S1 for complete table).

Identified behaviour components (n =164) were titled using accepted terminology and descriptions, published in Hanlon and Messenger^16^ and peer reviewed articles^15,26,42–44^. Terminology defining analogous behaviours witnessed in distantly related Coleoid families, were also incorporated into ethograms. Some pre-existing behavioural terms were manipulated to accommodate similar undescribed behaviours while logical labels were invented for non-reported behaviours (see Supplementary Table S1 & S10).

Ethograms were catalogued into Behavioural Observation Research Interactive Software (BORIS, v. 7.4)^45^. Behaviours which displayed a plethora of colour variations, e.g. mantle-margin stripe, were not distinguished as unique, to avoid extensively long ethograms (Supplementary Table S1).

### Behavioural recording

Non-mutually exclusive point (quick single behaviours) and state events (persistent behaviours) were categorised and recorded continuously (upon exhibition) throughout each feeding event. Inconsistently observed chromatic behaviours displayed on the cornea or ventral mantle (below fin) were not recorded.

Prey items were identified to general taxonomic group of Osteichthyes (bony fish) or Crustacea (e.g. crabs, shrimp), to eliminate species-level identification errors. Their respective benthic or pelagic lifestyles were determined using Debelius^46^ and Campbell^47^.

We removed behaviours thought to have no relation with hunting and grouped some well-understood and accepted body patterns and behaviours, e.g. mottled body pattern and behaviours that appear objectively like others – e.g. “downward curled arms” and “drooping arms” (see Supplementary Table S1). We then collated conspecific behavioural strings into transposed rows for Behatrix – Behavioural Strings Analysis v. 0.4.4^48^. These behavioural strings were then used in conjunction with Graphviz v. 2.38 ‘dot’^49^ package to visually depict transition(s) from one behavioural unit to the proceeding, with percentage values of relative occurrences equating to one^50,51^. Permutation tests were computed on transition matrices, based on observed counts of behavioural transitions, via Behatrix ‘Run random permutation test’ function. Each species dataset was permuted 100,000 times (shuffling data 100,000 times achieves: p = 0.05 ± 0.1% uncertainty, minimum value p = 0.00001), ranking the real test statistic amid shuffled test statistics and attaining p-values for each unique behavioural transition^51,52^.

### Correlation Tests

To provide evolutionary relationships from feeding behaviours, a dendrogram representing species relationships by shared behavioural units was constructed using presence/absence of behavioural units amongst all species. Binary data was bootstrapped/resampled 100 times using PHYLIP ‘SEQBOOT’ routine^53^. Resampled data was entered into ‘MIX’ routine for (multiple) tree construction by Wagner’s parsimony, which considers underlying ancestral states. Diverse dendrograms were combined into one using ‘CONSENSE’ routine. NEWICK.txt file output dendrogram with Figtree v. 1.4.4^54^.

We performed a conventional R-mode principal components analysis (PCA)^55^, on the primary data set of 156 behaviours observed in 86 cuttlefish (24 *M. pfefferi*, 13 *S. apama*, 8 S. *latimanus*, 17 *S. officinalis* and 23 *S. pharaonis*) via the Multivariate Statistical Package (MVSP)^56^.The data are dominated by null entries, with only 16% ‘1’-scores, hence the initial PCA extracted only 24.9% of available variance from axes 1 and 2 with the remaining 75.1% scattered across axes 3 to 84. To counter this, we removed single occurrence and then ‘rare’ (n ≤ 5) behaviours raising ‘1’-scores to 19.6% and 28.1% respectively with minimal impact on PCA axis 1/2 plot structure. While successful, this highlighted that most behaviours are restricted to a few, usually conspecific, individuals and so conventional two-axis PCA will only show a small fraction of the inherent signal. To counter this we imported PCA case scores and respective axial variance extraction (as a percentage of total) to MS Excel, where we multiplied case scores by percent variance and then summed ‘odd’ versus ‘even’ axes. We returned the data to MVSP and presented them as a two-axis PCA (i.e. linear regression). Graph topology is very similar to the conventional Axis 2/ Axis 1 original but contains all available signal. The odd/ even ‘stacking’ proved more successful than both ‘axis 1 versus 2-84’ or ‘top 50% versus bottom 50%’ in terms of similar topology to the ‘axis 2/ axis 1 original’.

## Results

A total of 85 feeding events were analysed, all included prey seizure attempts, and 63 resulted in successful prey seizure: *M. pfefferi* (20/24), *S. apama* (10/13), *S. latimanus* (5/8), *S. officinalis* (12/17), and *S. pharaonis* (16/23) (Table 2). Some 63 observations were concluded by a whole – or part (i.e. head or mantle section) – body chromatic pattern ‘flash’ display, after predation attempt. This behaviour was performed by all species (Table 2).

A total of 164 unique behaviours were recorded from all feeding events analysed in this study (see Supplementary Table S1): (3) “textural”, (7) “postural (whole body)”, (17) “postural (arms)”, (26) “locomotor”, (98) “chromatic”, and (13) “excluded behaviours”– all behaviours excluded from kinematic diagrams; only “freeze”, “clouding prey” and “unintentional prey startle” were removed from Wagner’s parsimony and PCA. Chromatic behaviours were either dynamic^20^ (17) or static (81), and specific displays varied in colour and physical attribute (surface area). The five species studied here shared 24 (15.09%) behaviours in total, and *Sepia spp*. shared 38 (23.90%) behaviours – both statistics include “excluded behaviours” (see Supplementary Table S1). Behaviours shared by heterospecifics were not all identical in form. For example, *M. pfefferi* achieved “tripod” posture with four points of benthic contact (e.g. fourth arm pair and two posterior ventral mantle “*glutapods*”^26^), whereas *Sepia spp*. supported themselves at three points (e.g. fourth arm pair and posterior ventral mantle).

### Prey types selection preferences and hunting modes

Cuttlefish species hunted specific prey types. Pelagic Osteichthyes – bony fish – were attacked more than benthic bony fish by *S. apama* (75%) and *S. pharaonis* (86.7%) (Table 3a), whilst *S. latimanus* equally preyed-upon both benthic (50%) and pelagic (50%) Osteichthyes (Figure 2B). All studied species, except *S. apama*, which was not observed hunting crustacea at all, targeted benthic Crustacea (100%) only (Table 3a). Despite proving harder to catch by all predators, a non-significant strong trend shows Osteichthyes (64.9% avg. predation success) were targeted more than crustaceans (90% avg. predation success) (Table 3B & Figure 2C). *S. latimanus* was unsuccessful at capturing bony fish (50% predation success) (Table 3b); and *S. officinalis* (60% predation success) was the only species that failed when predating crustaceans (Table 3b). *S. apama*, and *S. pharaonis*, had the least heterogenic diets of the five species, significantly selecting (Chi-square: X^2^_1_= 12, N = 12, p < 0.001, and X^2^_1_= 8, N = 18, p = 0.005: respectively) towards Osteichthyes prey. *M. pfefferi’* diet was the most varied (Chi-square: X^2^_1_= 1.19, N = 21, p = 0.275) (Table 3C & Supplementary Figure S2).

**Figure 2.**
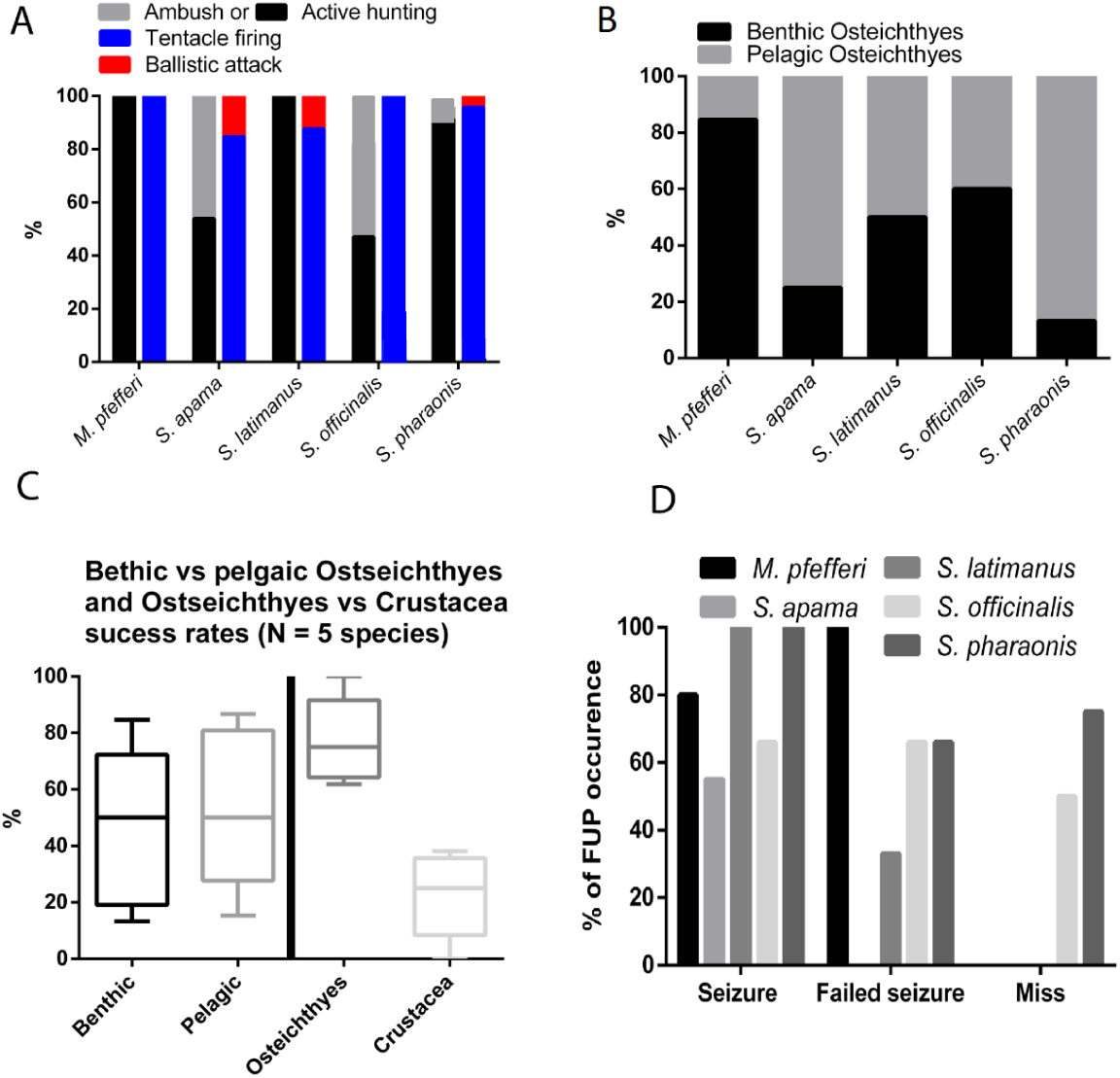
Differences in prey chosen, strategy and success were found in five cuttlefish species when taking observations from citizen science and social media: (A) Hunting strategy; (B) proportion of success rate in habitat occupancy in Osteichthyes; (C) Mean success rate, from proportions in habitat occupancy when all five species are combined (Benthic vs pelagic Wilcoxon Matched pairs P = 0.875) and also by prey type (Wilcoxon matched pairs P = 0.062); and lastly (D), per cent of ‘Flash upon predation’ (FUP) after successful prey seizure, failed seizure and complete miss. Five species include *M. pfefferi* (N = 24), *S. apama* (N = 13), *S. latimanus* (N = 8), *S. officinalis* (N = 17), and *S. pharaonis* (N = 23).

*S. apama, S. officinalis*, and *S. pharaonis*; captured prey via ambush, and active hunting strategies, whereas *M. pfefferi* and *S. latimanus* obligately hunted actively (Figure 2A). *S. officinalis* and *S. pharaonis* executed both active, and ambush modes of hunting, during single observations (Figure 2A). Prey was caught via tentacular firing, or tactile arms – when ballistic attacks were employed. *Sepia spp*. practiced both methods of prey capture, whilst *M. pfefferi* displayed no deviation from “tentacle firing” (Figure 2A).

### Kinematic diagrams

Significant probabilities (p < 0.05 and p < 0.01) for behavioural transition occurrence are represented as conspecific kinematic diagrams in thin vs thick black lines (Figure 3). Figure 3 shows that all studied species incorporate at least one, if not multiple types of conspicuous chromatic signals (signals: pursuit-deterrent signal – “flash upon predation”; aposematic display “warning”, and dynamic skin pattern – “flash”, “chromatic pulse” and “rhythmic passing waves”), during hunting – details of such conspicuous displays are described below and see Supplementary Table S1 for ethogram and behaviour descriptions.

**Figure 3.**
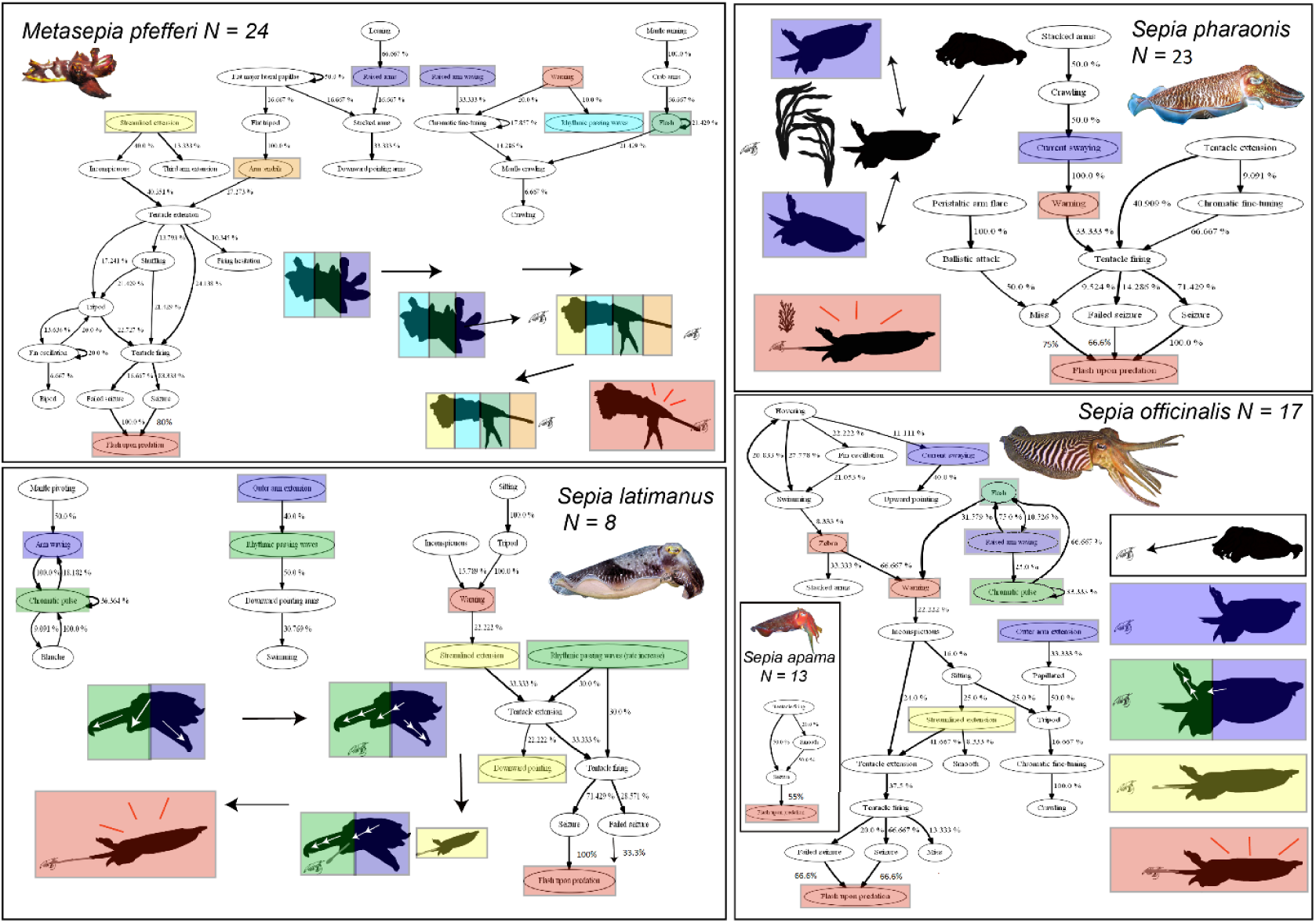
A selection, for brevity, of significant (p = 0.05 (thin lines) – p < 00.1 (thick lines)) kinematic diagrams for the five species of cuttlefish studied here: e.g. *M. pfefferi* (N = 24), *S. apama* (N = 13), *S. latimanus* (N = 8), *S. officinalis* (N = 17), and *S. pharaonis* (N = 23) from data gathered via retroactive citizen science requests or social media. See Supplementary Figure S3-S7 for all significant chains for all species: Note, all (one) significant transitions for *S. apama* are included here due to there being so few. Coloured boxes highlight different signals employed during hunting, including physical/postural changes (i.e. streamlined extension, arm raising/waving), chromatic (warnings, flashes, pules and dynamic displays) and ‘flash upon predation’.

- *M. pfefferi*: “Rhythmic passing waves”, flashing (21.43%, p = 0.01), “warning”, and aposematic display adjustments (i.e. “chromatic fine-tuning) (17.86%, p = 0.05), were exhibited during foraging and positioning stages, prior to “tentacle firing”. “Flash upon predation” was signalled after both successful (80%, p < 0.001) and unsuccessful (100%, p < 0.001) prey seizure attempts.
- *S. apama*: Showed “flash upon predation” after successful prey capture (55.56%, p < 0.001), only – “seizure” also transitioned to “smooth” (20%, p = 0.32) textural component.
- *S. latimanus*: “Warning”, “rhythmic passing waves” and recurring “chromatic pulse” (36.36%, p < 0.001) displays were present during foraging and positioning periods. The rate of “rhythmic passing waves” increased before both, extension (30%, p = 0.005) and firing (30%, p = 0.002) of feeding tentacles. Only successful prey seizure was significantly reciprocated by “flash upon predation” (100%, p < 0.001).
- *S. officinalis*: Foraging and positioning events included chromatic pulsing (33.33%, p = 0.019), “flash”, and “warning” signals, such as “zebra” displays. Pursuit-deterrent signal was displayed upon both, “seizure” (66.66%, p < 0.001) and “failed seizure” (66.66%, p= 0.003).
- *S. pharaonis*: Repetitive “flash” (66.67%, p < 0.001) displays and aposematic body patterns preceded tentacle firing (33.33%); and “seizure” (100%, p < 0.001), “failed seizure” (66.66%, p = 0.012) and “miss” (75%, p < 0.001) were all concluded with “flash upon predation”.

The cuttlefish species studied here used directive marks, by performing dynamic postural mechanisms (i.e. “arm waving”) alone, or in combination with dynamic skin patterns: *M. pfefferi* repeatedly twitched raised arms (66.67%, p < 0.001) and manipulated “arm tendrils” (27.27%, p = 0.04) before “tentacle extension”; *S. latimanus* operated “arm waving” with “chromatic pulse”; and *S. officinalis* either combined “flash” with “raised arm waving”, or extended this routine by incorporating “chromatic pulse” (66.67%, p = 0.01) before “flash”, and after “raised arm waving” (25%, p = 0.04).

### PCA

Covariance of behavioural units separates all studied *Sepia* spp. (negative x-axis) from *M. pfefferi* (positive x-axis) (Figure 4A). *S. officinalis* and *S. pharaonis* have observations present in all four quadrants, and that overlap with all other *Sepia* spp. (Fig. 4B); whereas *S. apama* and *S. latimanus* observations show a more precise spread, and do not overlap with each other (Figure 4A/B).

**Figure 4.**
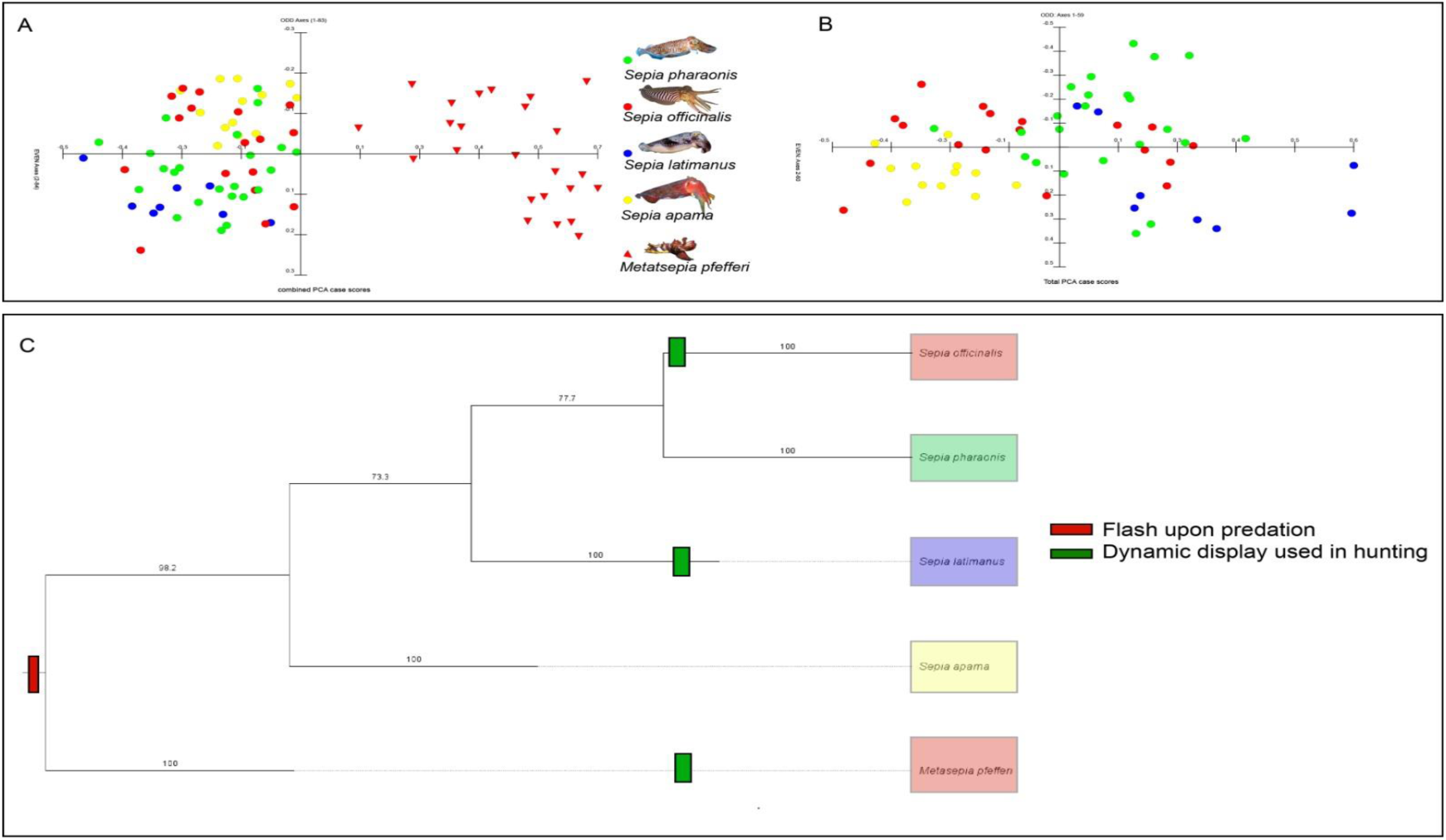
*M. pfefferi* (N = 24), *S. apama* (N = 13), *S. latimanus* (N = 8), *S. officinalis* (N = 17), and *S. pharaonis* (N = 23) shared behaviours gathered from social media and retroactive citizen science requests: PCA all studied species (A); PCA *Sepia* spp. only (B); and Wagner’s parsimony and conserved behaviours of interest: pursuit-deterrent signal (i.e. “flash upon predation” and dynamic skin patterns (i.e. “chromatic pulse” and “rhythmic passing waves”) (C)

### Correlations of ‘behavioural traits’ (Wagner’s parsimony)

Wagner’s parsimony concluded confident (73.3% – 100%) branching of species studied, based on shared behaviours (Figure 4C) (see Supplementary Table S1). *M. pfefferi* is grouped separate (100%) from, but more relative to *Sepia* spp. (98.2%). *S. apama* shares the least vestigial behaviours with other *Sepia* spp., and *S. officinalis* and *S. pharaonis* are most related (77.7%), based on shared behaviours (Figure 4C).

## Discussion

Cuttlefish prey selection appeared opportunistic – each species targeted various crustacean or bony fish species and target selection could have been influenced by the variability of seasonal prey availability^57^. However, we saw a very strong, albeit non-significant trend (Figure 2) towards a predominant Osteichthyes diet. We suspect that increasing the species (i.e. N > 5 used here) number would produce a significant difference between crustacea and bony fish for adult cuttlefish and is expected amongst mature cephalopods^13,57^ like the ones studied here. Adults are not obliged to crustacean only diets, unlike juveniles (crustacea are rich in amino acids and polypeptides, essential nutrients for juvenile cephalopod growth^13,57^), and a varied fish dominant diet becomes of greater nutritional value into adult life^57–59^.

The species studied here used different hunting strategies (Table 1) during feeding events (Figure 2 & 3). Tactics ranged from ambush (e.g. *S. apama, S. officinalis*, and *S. pharaonis*) to mobile: prey-stalking, accelerated prey-pursuit, and speculative trapping; collectively referred to as “active hunting”. Cuttlefish of all species were seen transitioning between distinct hunting strategies within single observations. Comparable strategy cycling has been witnessed amongst other predatory taxa, like Salticids (jumping spiders), and is recognised as a co-evolutionary mechanism that conditionally reciprocates to real-time prey behaviour^60,61^, enabling predators to overcome situational predatory challenges such as environmental obstacles or prey awareness^60^.

Both S. *officinalis* and *S. apama* showed ∼50% ambush hunting (Figure 2) which is far higher than the other three species (∼0-10%). This is may be explained by two different reasons. *S. officinalis* are found in the coolest waters of all the cuttlefish here and maybe be conserving energy for growth; their water is also likely the most turbid (Cooke pers.obs) suggesting an opportunistic approach may be better in poor visibility conditions. *S. apama* however, are found in more tropical waters like the remaining three species but are thought to be the world’s largest cuttlefish^62^ and so may again be conserving energy in movement, this time for growth. Why they are the biggest is presently a mystery but may therefore have some selective advantage if they are actively avoiding energy expenditure by using ambush hunting at least as much as active hunting.

Tentacular firing was preferentially used over ballistic attack to capture both crustaceans and fishes (Figure 2). Rapid ejection (30-75 ms)^63^ of feeding tentacles achieves high predation accuracy by supressing prey reaction time^16,63^. We observed cuttlefish altering directions of gap-closing pathways (e.g. *S. latimanus, S. officinalis*, and *S. pharaonis*) and performing other transient ballistic attack tunings (see Supplementary Figure S5, S6, & S7).

Directive marks (e.g. *M. pfefferi, S. latimanus*, and *S. officinalis*) and mimicry (e.g. *M. pfefferi*, and all *Sepia spp*. looked at here) was observed within active hunting or ambush events, suggesting that they are not independent hunting modes themselves, as previously thought^16^, but rather body patterns (chromatic, textural, postural) or locomotor components that can be simultaneously/sequentially expressed and aid hunting by the addition of prey distraction or prey deception^20,21,26^.

Cuttlefish, and other Coleoid cephalopods, possess a very large number of quantifiable behaviours when posture, chromatic and locomotor behaviours are all included, compared to possibly any other taxa, and we can therefore theoretically derive evolutionary relationships of species in such manner analogous to morphological and molecular taxonomy, using the presence or absence of specific behavioural traits^64,65^. Despite having prey selection differences^63,66,67^ (Fig. 2) we found hunting behaviour was most similar between *S. officinalis* and *S. pharaonis* (Fig. 4), which makes sense given their close phylogenetic relationship^28,68^ and bordering range proximity (e.g. SE Mediterranean Sea – Gulf of Suez/Red Sea)^69–71^. However, the majority of our observations were conveyed on geographically distant *S. pharaonis* that inhabited Indo-Pacific waters (Fig. 1); and observations potentially included a different sub-species of *S. pharaonis*, recent molecular studies reveal the species may be a species complex – consisting of three to five sub-species^69,70^. Localised resource partitioning^72^, or seasonal changes in hormones^73^ not controlled for here, or divergent prey selection^74^might have caused the divergence in hunting behaviour^73,74^.

Kinematic diagrams (Figure 3) show non-random behavioural transitions, with respective probabilities of transitional occurrence from preceding behaviour; core hunting components for species studied here – significant transitions can have small percentage values when preceding behaviours lead into many different behaviours. Species kinematic diagrams vary in behaviours, behavioural transitions, and transitional chain-lengths. Short (i.e. subjective length) behavioural sequences suggest less behaviours were expressed during our observations^75,76^, or a product of high transitional variability^76^ – this may explain why certain sequences terminate before “tentacle firing” (Figure 3). *S. apama*’s hunting behaviour was least modal and uniquely abstained from conspicuous displays (e.g. dynamic skin patterns, directive marks), and even deimatic (i.e. warning) displays. However, they also had the shortest average videos, and showed the fewest overall behaviours. They also used the fewest ‘flash upon predation’ than the other species studied here. *S. apama*, or the “Giant Australian Cuttlefish”, can produce extremely conspicuous dynamic displays during courtship^20,77^, but for unknown reasons seem to not use them during hunting in the videos we analysed (N = 13), perhaps being so large has reduced the possible number of predators.

Being both predator and prey is a significant factor for cephalopod behaviour^2^. Subduing, killing, via protein-based neurotoxins^13,78^ and assimilating prey is not an immediate process^11^. It can take time, and this makes cuttlefish vulnerable to kleptoparasitism and predation from other predators that can detect and trace signs of an animal being killed and or eaten (i.e. vibration, blood etc)^11,79^. To deal with this threat the species studied here appear to use a ‘pursuit-deterrent signal’^7–9^ which we call “flash upon predation”. Its function may startle or deter predators/competitors during vulnerable prey handling periods by showing possible awareness to (even if the cuttlefish is not actually aware of a predator nearby) or fitness^11^. Despite not having observed any prey handling cuttlefish becoming a victim of heterospecific competition/kleptoparastism, nor witnessing any conspecifics present in any videos (except *M. pfefferi -* see Supplementary Table S10), we suggest that being eaten by one of their many predators is a pressing concern for distracted cuttlefish. “Flash upon predation” was selectively reciprocated to “tentacle firing” outcomes (i.e. successful “seizure”, “failed seizure”, and sometimes even prey “miss”), suggesting its use is situational rather than obligatory. We cannot rule out that the signal was directed at the SCUBA diver/s but its function would likely still be the same – a warning of some kind. However, generally, almost all cuttlefish encountered by divers appear to be more or oblivious to their presence unless followed very closely. They will nearly always continue what they are motivated to do (Cooke pers.obs). We did not include videos in our analysis where the divers appeared to alter the behaviour of a cuttlefish. Like other relative flashing patterns observed in this study, “flash upon predation” varied between species and individuals within species (Fig. 2), some displays were more conspicuous (contrasting more with predominant body colour) and or occurred over larger body areas.

There are many examples in the animal kingdom of the use of pursuit-deterrent signals, termed “flash upon predation” here, towards predators as a form of defence; these signals can be highly variant across taxa, but sometimes show convergent aspects in form and function: Polynoids and Ophiuroids eject sacrificial luminescent lures^80^; collector sea urchins (*Tripneustes gratilla*) release clouds of venomous pedicellaria heads^81^; New World tarantulas (Theraphosidae) brush off venomous barbed abdominal or palpal urticating hairs (only investigated when being predated)^82,83^; bombardier beetles (*Metrius contractus*) spray hot quinonoid secretions^84^. Other species rely on signals, chipmunks (*Tamias striatus*) and grey squirrels (*Sciurus carolinensis*) harass timber rattlesnakes (*Crotalus horridus*) by repeatedly approaching them whilst tail-flagging^85^. As far as we know our study is the first study to show pursuit deterrent signals in complex marine predators but also, perhaps more interestingly, in species that are both predator and prey.

## Conclusions

This is the first study of its kind that uses social media and citizen science to investigate important behavioural ecology processes that add to animal hunting behaviour theory (i.e. pursuit deterrent signals in a predator, which is also prey). We also employ novel analysis (phylogeny’s based on behavioural sequences) that may help resolve taxa where typical methods have failed in cuttlefish. Lastly, we show that cuttlefish use a mixture of warning and predatory behaviours throughout hunting sequences. Whilst limitations exist, as discussed, we believe the untapped resource of unsolicited animal behaviour observations from social media may provide valuable knowledge at minimal economical cost.

