## Supplementary material for "Social Media and citizen science provide valuable data for behavioural ecology research: Are cuttlefish using pursuit-deterrent signals during hunting?"

*Corresponding author

Supplementary materials “Combined species ethogram”

Supplementary Table S1. All displayed behaviours (species: *M. pfefferi*, *S. apama*, *S. latimanus*, *S. officinalis*, and *S. pharaonis*) assorted into components: Texture, Postural (whole body), Postural (arms), Locomotor, Chromatic, and Excluded behaviours. Component titles are positioned in the column furthest to the left, with constituent behaviours indented into less marginal columns. ‘Excluded behaviours’ were not incorporated into Markov chain analysis, to reduce complexity of output sequences; and due to rare frequency. Behaviours were unevenly assorted between species: *M. pfefferi* = ■, *S. apama* = ▲, *S. latimanus* = ►, *S. officinalis* = ▼, *S. pharaonis* = ◄.

| **Behaviours and components** | | **Behavioural definitions** | **Species displayed** |
|---|---|---|---|
| **Texture** | | | |
| 1 | Flat major lateral papillae | Major lateral papillae are relaxed, not prominent | ■ |
| 2 | Papillated | Expressed three-dimensional skin texture (can be displayed on head, arms, or mantle only) | ■ ▲ ► ▼ ◄ |
| 3 | Smooth | Fully relaxed smooth skin texture (can be displayed in isolation on head, arms, or mantle only) | ■ ▲ ► ▼ ◄ |
| **Postural (whole body)** | | | |
| 4 | Bipod | Cuttlefish balances on its fourth pair of arms, whilst mantle is elevated from substrate | ■ ▲ ► ▼ ◄ |
| 5 | Downward pointing | Straight body profile orientated downwards | ▲ ► ▼ ◄ |
| 6 | Flat tripod | Fourth pair of arms are completely extended, head and ventral portion of mantle are in complete contact with substrate | ■ |
| 7 | Leaning | Centre of gravity is positioned over one anchored fourth arm | ■ |
| 8 | Sitting | Cuttlefish is negatively buoyant and laying on substrate | ▲ ► ▼ ◄ |
| 9 | Tripod | Fourth pair of arms and mantle pseudopods resemble a tripod like stance, from which the feeding tentacles are fired | ■ ► ▼ ◄ |
| 10 | Upward pointing | Straight body profile orientated upwards | ▲ ► ▼ ◄ |

| | | | |
|---|---|---|---|
| **Postural (arms)** | | | |
| 11 | Crab arms | Arms possibly mimic that of a crab's claws and antenna | ■ |
| 12 | **Downward pointing arms** | Constituted by drooping arms, downward curled arms, and lagging fourth pair | ■ ▲ ▶ ▼ ◀ |
| 13 | Downward curled arms | Constituted by drooping arms, downward curled arms, and lagging fourth pair | ▶ ▼ ◀ |
| 14 | Drooping arms | Arms are orientated downwards with tips curled towards posterior end | ■ ▲ ▶ ▼ ◀ |
| 15 | Lagging fourth pair | Bulk of arms are orientated downwards | ▲ ▶ ▼ ◀ |
| 16 | Forward jetting arms | Arms are orientated towards posterior end due to high speed swimming | ◀ |
| 17 | **Outer arm extension** | Fourth or third arm pair(s) are splayed at a 45-degree angle from central arm cluster | ▲ ▶ ▼ |
| 18 | Third arm extension | Third arms splayed at a 45-degree angle from central arm cluster. Fourth arm pair are planted on substrate and used to crawl | ■ |
| 19 | Peristaltic arm flare | Arms are constricted at the base whilst ends are flared | ▲ ▼ ◀ |
| 20 | **Raised arms** | The first (or both first and second) pair of arms are raised upwards, noticeably separate from remaining arms. Raised arms are distinctly: fully erect, downward curling at tips, or crinkled | ■ ▲ ▶ ▼ ◀ |
| 21 | Raised arms (crinkled) | Raised arms characteristically crinkled in texture and shape | ▲ ▼ ◀ |
| 22 | Raised arms (curling tips) | Raised arms are characterised by curling tips | ▲ ▶ ▼ ◀ |
| 23 | Raised arms (net) | Raised arms broaden, increasing in width | ▲ |
| 24 | Raised arms (straight) | Raised arms are fully erect | ▲ ▶ ▼ |
| 25 | Rigid arms | Arms constricted and pointing directly ahead, extending from body axis | ■ ▼ |
| 26 | Stacked arms | Arms are stacked in corresponding pairs | ■ ▲ ▶ ▼ ◀ |
| 27 | Streamlined extension | Arms are pointing directly ahead, extending from body axis | ■ ▲ ▶ ▼ ◀ |
| **Locomotor** | | | |
| 28 | Arm tendrils | Wriggling tendril-like projections extend from the third arm pair | ■ |
| 29 | Arm waving | Arms (can be both first and second pair) abruptly raised and lowered in short repetitive succession | ▲ |

| | | | |
|---|---|---|---|
| 30 | Ballistic attack | Head on launch (at prey position) and seizure by arms only | ▲ ▶ ▼ ◀ |
| 31 | Ballistic attack hesitation | Head on launch (at prey position) is aborted before attempt of seizure | ▶ |
| 32 | Crawling | Fourth pair of arms used to crawl/ walk along the benthos or solid environmental objects | ■ ▲ ▶ ▼ ◀ |
| 33 | Current swaying | Cuttlefish sways with water current, mimicking algal or soft coral movement | ▲ ▶ ▼ ◀ |
| 34 | Drooping arms ripple | Drooping arms are motioned in a whip-like fashion | ◀ |
| 35 | Failed seizure | Prey item is unsuccessfully secured by feeding tentacles | ■ ▲ ▶ ▼ ◀ |
| 36 | Fin oscillation | Kinetic stimulation of radial mantle fin | ■ ▲ ▶ ▼ ◀ |
| 37 | Firing hesitation | Extended feeding tentacles are retracted before their firing | ■ ▶ ▼ ◀ |
| 38 | Flight (jet) | Siphon ejects water causing a backwards burst | ▼ |
| 39 | Hovering | Cuttlefish remains stationary in the water column | ▲ ▶ ▼ ◀ |
| 40 | Mantle crawling | Fourth pair of arms and mantle glutapods used to crawl/walk on environmental substrate | ■ |
| 41 | Mantle pivoting | Buoyancy of cephalic region is increased and decreased to achieve pivoting | ▶ |
| 42 | Mantle running | Fourth pair of arms and mantle glutapods used to run along environmental substrate | ■ |
| 43 | Miss | Prey item is missed on attempt of seizure | ▲ ▼ ◀ |
| 44 | Prey circling | Detected prey is circled with head on attention | ▲ ▶ ◀ |
| 45 | Raised arm twitching | Raised arms are repetitively twitched | ■ |
| 46 | Raised arm waving | Raised arms are waved upwards and downwards | ■ ▼ ◀ |
| 47 | Sediment burial | Cuttlefish buries itself in fine sediment | ▼ |
| 48 | Sediment probing | Arms probe downwards into fine sediment | ▶ |
| 49 | Seizure | The prey item is seized by either: firing feeding tentacles or tactile arms | ■ ▲ ▶ ▼ ◀ |
| 50 | Shuffling | Cuttlefish shuffles side to side on benthos | ■ |
| 51 | Swimming | Cuttlefish buoyantly locomotes using fin oscillation for propulsion | ▲ ▶ ▼ ◀ |
| 52 | Tentacle extension | Feeding tentacles are extended | ■ ▲ ▶ ▼ ◀ |

Pre-print

| | | | |
|---|---|---|---|
| 53 | Tentacle firing | Feeding tentacles are fired | ■ ▲ ▶ ▼ ◀ |
| **Chromatic** | | | |
| 54 | Blanche | Entirely white in appearance | ▶ ▼ ◀ |
| 55 | **Chromatic fine tuning** | Dynamic displays dissociative of passing cloud behaviour | ■ ▲ ▶ ▼ ◀ |
| 56 | Darkening | Pigmentation darkens in colouration | ▼ |
| 57 | Fin stripe (colour change) | Colour of fin stripe changes | ▶ |
| 58 | Mantle margin stripe (colour change) | Colour of mantle margin stripe changes | ▶ |
| 59 | Ocular epaulettes (colour change) | Colour of ocular epaulette changes | ■ |
| 60 | Paired head spots (colour change) | Colour of paired head spots changes | ■ |
| 61 | Paling | Overall paling in colouration | ■ ▲ ◀ |
| 62 | Papillae outline (colour change) | Colour outlining major lateral papillae changes | ■ |
| 63 | Raised arms (colour change) | Raised arms change in colouration | ▶ |
| 64 | Walking arm spots (colour change) | Walking arm (fourth pair) spots change in colouration | ■ |
| 65 | Walking arms (colour change) | Walking arms change in colouration | ■ |
| 66 | Chromatic pulse | Wandering flash display | ▶ ▼ |
| 67 | Flash | Brightening or abrupt change of localised pigmentation | ■ ▶ ▼ ◀ |
| 68 | Flash upon predation (pursuit-deterrent signal) | Brightening or abrupt change of pigmentation upon predation attempt | ■ ▲ ▶ ▼ ◀ |
| 69 | **Inconspicuous** | Chromatic displays that do not attract attention | ■ ▲ ▶ ▼ ◀ |
| 70 | Crab head | Pigmentation resemblant of a crab (orange, black, yellow) | ■ |
| 71 | **Disruptive** | Highly contrasting dark and light patches assembled in a non-repetitive configuration | ▼ ◀ |
| 72 | Animation eyes | White eye patch with a black outline | ■ |
| 73 | Anterior head stripe | Stripe between eyes. Often bordering head bar | ▶ |

| | | | |
|---|---|---|---|
| 74 | Anterior mantle patch | Pigmented patch displayed on the anteriorly on the mantle | ▼ |
| 75 | Arm bands | Thin lateral Stripes lining the arms | ■ ▲ ▶ ▼ ◀ |
| 76 | Banded arms | Thick lateral Stripes lining the arms | ▶ |
| 77 | Banded by lines (transverse) | Transversely banded mantle by wave-like anterior and posterior mantle lines. Lines segregate mantle pigmentation into contrasting (light/dark) blocks | ▲ ▶ ▼ ◀ |
| 78 | Black arm tips | Ends of arms are black coloured | ■ |
| 79 | Clear fins | Fins are devoid of colour | ■ ▲ ▶ ▼ ◀ |
| 80 | Dark arms | Arms are dark in colouration in comparison to predominant body colour | ▶ ▼ ◀ |
| 81 | Dark head pale arms | Head is of dark pigmentation in comparison to arms | ■ ▲ ▼ ◀ |
| 82 | Darker mantle | Mantle is of darker colouration than head and arms | ■ ▲ |
| 83 | Dorsal mantle bars | Pigmented bars orientated perpendicular to mantle margin | ■ |
| 84 | Dorsal mantle bars (spots) | Pigmented bars with spots orientated perpendicular to mantle margin | ■ |
| 85 | Dorsal stripe | Stripe/band ranging centrally from anterior to posterior region of the mantle | ■ ▼ ◀ |
| 86 | Flamboyant | Highly disruptive body pattern composed often accompanied by dark brown, white and yellow pigmentation | ■ |
| 87 | Flamboyant head | Flamboyant pattern on head only | ■ |
| 88 | Fragmented arms | Arms are fragmented by dark patches | ▶ |
| 89 | Head bar | Pigmented bar displayed on the top section of the head, posteriorly to eyes | ▶ ▼ ◀ |
| 90 | Helmet patch | Pigmented bar patch with a perpendicular protrusion between the eyes | ▲ |
| 91 | Mantle square | Pigmented square displayed centrally on mantle | ▼ ◀ |
| 92 | Median mantle stripe | Opposingly coloured stripes border central dorsal stripe | ◀ |
| 93 | Pale arms | Arms are pale or clear (absent of pigmentation) | ◀ |
| 94 | Pale fourth pair | Fourth pair of arms are white and pattern-less | ■ |

Pre-print

| | | | |
|---|---|---|---|
| 95 | Pigmented tentacular clubs | Feeding tentacles are not uniformly pale, clubs are coloured | ■▲▶▼◀ |
| 96 | Zebra face | Static vertical black bands overlay predominantly white head pigmentation | ■ |
| 97 | **Mottled** | A pattern characterised by small or large dark or light oval-shaped patches that are repeated across the animal’s body | ■▲▶▼◀ |
| 98 | Arm spots (dark) | Small dark spots lining arms | ▲▶▼◀ |
| 99 | Arm spots (white) | Small white spots lining arms | ▲▶▼◀ |
| 100 | Broken mantle margin stripe | Mantle margin stripe is discontinuous | ■ |
| 101 | Mottled dorsal mantle | Mottle pattern is present on central portion of mantle only | ▶ |
| 102 | Pigmented papillae | Papillae are of different pigmentation to predominant colour | ▲ |
| 103 | Radial mantle stripes | Dark lines (long or short) extending perpendicularly from mantle margin and towards mantle centre. This pattern is displayed along the entire perimeter of the mantle | ▶▼◀ |
| 104 | Small spots (dark) | Body is littered with small dark spots | ■▲▶▼◀ |
| 105 | Small spots (white) | Body is littered with small white spots | ■▲▶▼◀ |
| 106 | **Uniformly stippled** | A uniform body colouration displaying minimal contrast in overall colour (accompanied by small dark or light patches/ spots) | ■▲▶▼◀ |
| 107 | Dark head and arms | Head and arms are darkly pigmented in comparison to mantle | ■▲▶▼◀ |
| 108 | Pale head and arms | Head and arms are pale in pigmentation in comparison to mantle | ■▶◀ |
| 109 | All dark | Entire body represents dark pigmentation, devoid of patterns | ■▼ |
| 110 | All purple | Body pigmentation is predominantly purple all over. Small white spots or yellow papulations may be present | ■ |
| 111 | All red | Body pigmentation is predominantly red all over. Small white spots may be present | ■ |

| | | | |
|---|---|---|---|
| 112 | **Rhythmic passing waves** | Dark bands on mantle travel from anterior mantle margin to posterior end, passing over static patterns | ■ ▶ |
| 113 | Converging rhythmic passing waves | Dark bands traveling from mantle posterior end and anterior margin converge at centre of mantle. Dark bands also travel away from central mantle point towards mantle margin flanks | ■ |
| 114 | Rhythmic passing waves (decrease rate) | The frequency of passing cloud display decreases | ▶ |
| 115 | Rhythmic passing waves (increase rate) | The frequency of passing cloud display increases | ▶ |
| 116 | **Warning** | Aposematic chromatic displays | ■ ▲ ▶ ▼ ◀ |
| 117 | Arm spots (dark) | First three pairs of arms are lined with large black spots | ■ |
| 118 | Arm spots (white) | First three pairs of arms are lined with large white spots | ■ |
| 119 | Arm stripe | A stripe outlining one side of an arm | ■ ▲ ▶ ▼ ◀ |
| 120 | Arm stripe connected eye ring | Stripe outlining one side of an arm extends to encircle the eye | ▶ ▼ |
| 121 | Crown | Crown-like patch displayed on posterior head boundary | ▲ |
| 122 | Central mantle spot | Large spot at the centre of the mantle | ◀ |
| 123 | Cheek patch | Pigmented patch displayed on either side of anterior head | ▶ ▼ ◀ |
| 124 | Rosy cheeks | Pink patch displayed on either side of anterior head | ▲ |
| 125 | Deimatic display | Paired dark mantle spots (false eye spots) accompanied with dark eye rings, dilated pupils, dark fin stripe, and general paling of body pigmentation | ▲ ▼ |
| 126 | Eye ring | Pigmented ring encircling the eyes | ▲ ▶ ▼ ◀ |
| 127 | Eye spot (behind) | Pigmented spot posterior to eyes | ▲ |
| 128 | Eyebrow patch | Pigmented patch above eyes | ▼ ◀ |
| 129 | Fin stripe | Line (coloured) on outer fin boundary | ▲ ▶ ▼ ◀ |
| 130 | Fourth pair blush | Large pigmented patch at top of arm | ■ |

| | | | | |
|---|---|---|---|---|
| 131 | | Fragmented dark mantle | Large pigmented patches arranged in circular formation around central mantle position | ▶ ◀ |
| 132 | | Infra-ocular patch | Pigmented patch in front of eyes | ■ ▶ ◀ |
| 133 | | Landmark spots | Four white spots present on top of head in an anterior leading arc-like formation. Central pair of spots are larger than peripheral pair | ◀ |
| 134 | | Large spots (dark) | Body is littered with large dark spots | ▲ ▶ ▼ ◀ |
| 135 | | Large spots (white) | Body is littered with large white spots | ▶ ▼ |
| 136 | | Mantle margin stripe | Stripe lining between the border between mantle and fin | ■ ▶ ▼ ◀ |
| 137 | | Ocular epaulettes | Block of colour or zebra-like pattern above, below, or completely encompassing eyes | ■ ▲ ▶ ▼ |
| 138 | | Paired arm spots | Medium sized dark spots on first pair of arms whilst raised | ▼ |
| 139 | | Paired head spots | Pair of spots on the posterior head | ■ ◀ |
| 140 | | Paired mantle spots | Pair of spots displayed centrally on the mantle | ■ ▲ |
| 141 | | Paired mantle spots (anterior) | Pair of spots displayed on the anterior portion of the mantle | ▼ |
| 142 | | Peppered fins | Fins are littered with small pigmented speckles | ▶ ▼ |
| 143 | | Pigmented major lateral papillae | Major lateral papillae are outlined by pigmentation | ■ |
| 144 | | Pink arms | First three pairs of arms are pink in coloration from their midpoint to tip | ■ |
| 145 | | Purple arms | First three pairs of arms are purple in coloration from their midpoint to tip | ■ |
| 146 | | Red arms | First three pairs of arms are red in coloration from their midpoint to tip | ■ |
| 147 | | Stitchwork fins | Disconnected mantle margin stripe (made from as sequence of spots) | ▼ |
| 148 | | Walking arm spots | Large spots lining the fourth pair of arms change colour | ■ |
| 149 | **Zebra** | | Zebra-like pattern chromatic display | ▼ ◀ |

| | | | |
|---|---|---|---|
| 150 | Strong zebra | A highly distinguishable zebra-like body pattern consisting of white and dark brown coloration. Can be selectively displayed on arms and head, or mantle only | ▼ |
| 151 | Weak zebra | A dull zebra-like body pattern consisting of white and light brown coloration. Can be selectively displayed on arms and head, or mantle only | ▼ ◀ |
| **Excluded behaviours** | | | |
| 152 | Attention | Upon detection of a prey item the cuttlefish may: change body patterning, erect first (and sometimes second) pair of arms, and align to face prey front on | ■ ▲ ▶ ▼ ◀ |
| 153 | Body twitch | Abrupt twitch or shake | ■ ▼ |
| 154 | Broad arms | All four pairs of arms are contracted and flattened to achieve broad appearance | ▶ ▼ ◀ |
| 155 | Clouding prey | Arms disturbed substrate, masking prey item | ▶ |
| 156 | Flattened | Entire body is widened, and eyes move from a lateral to upward position | ◀ |
| 157 | Freeze | Abrupt halt of all movement | ■ |
| 158 | On axis rotation | Cuttlefish rotates facing direction around central axis | ■ ▶ ▼ ◀ |
| 159 | Peripheral scanning | Head or eyes scan either side of head-on field of view | ■ ▶ ▼ ◀ |
| 160 | Raised dot | Tips of first pair are raised and darkly pigmented | ▼ |
| 161 | Shield | Ornamental patch at centre of mantle | ◀ |
| 162 | Smooth (head only) | Fully relaxed smooth skin texture (can be displayed in isolation on head only) | ■ ▲ ▶ ▼ ◀ |
| 163 | Turning | Cuttlefish changes direction whilst travelling | ■ ▲ ▶ ▼ ◀ |
| 164 | Unintentional prey startle | Prey item is physically startled by cuttlefish oblivious to its presence | ◀ |

#### Using social media and citizen science to drive behavioural ecology research: Are cuttlefish using pursuit deterrent signals during hunting?

Dražen Gordon, Philip Pugh and *Gavan M Cooke

Department of Life Sciences, Anglia Ruskin University, Cambridge, United Kingdom.

*Corresponding author

Supplementary materials “Predation success”

Supplementary Figure S2. Proportions of success attempts for two prey types – Osteichthyes (bony fish) or Crustacea

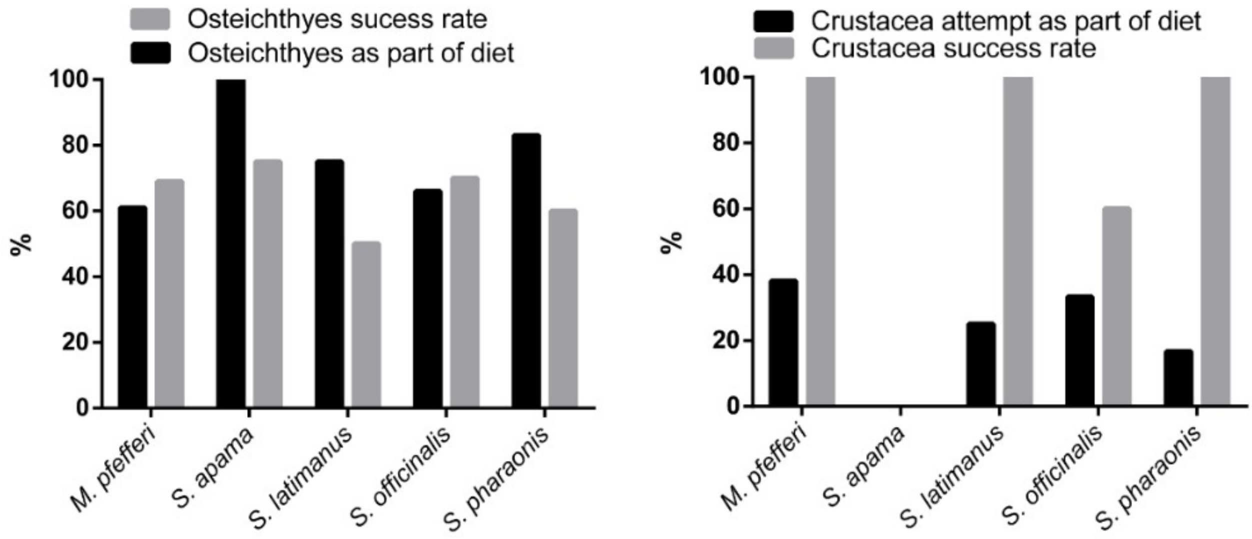


Using social media and citizen science to drive behavioural ecology research: Are cuttlefish using pursuit deterrent signals during hunting?

Dražen Gordon, Philip Pugh and *Gavan M Cooke

Department of Life Sciences, Anglia Ruskin University, Cambridge, United Kingdom.

*Corresponding author

Supplementary materials “Full kinematic figures”

Supplementary Figure S3. All behavioural transitions from all *Metasepia pfefferi* observations. Significant behavioural transitions are shown by line thickness ($p = 0.05$ (thick lines) – $p < 0.001$ (thicker lines))

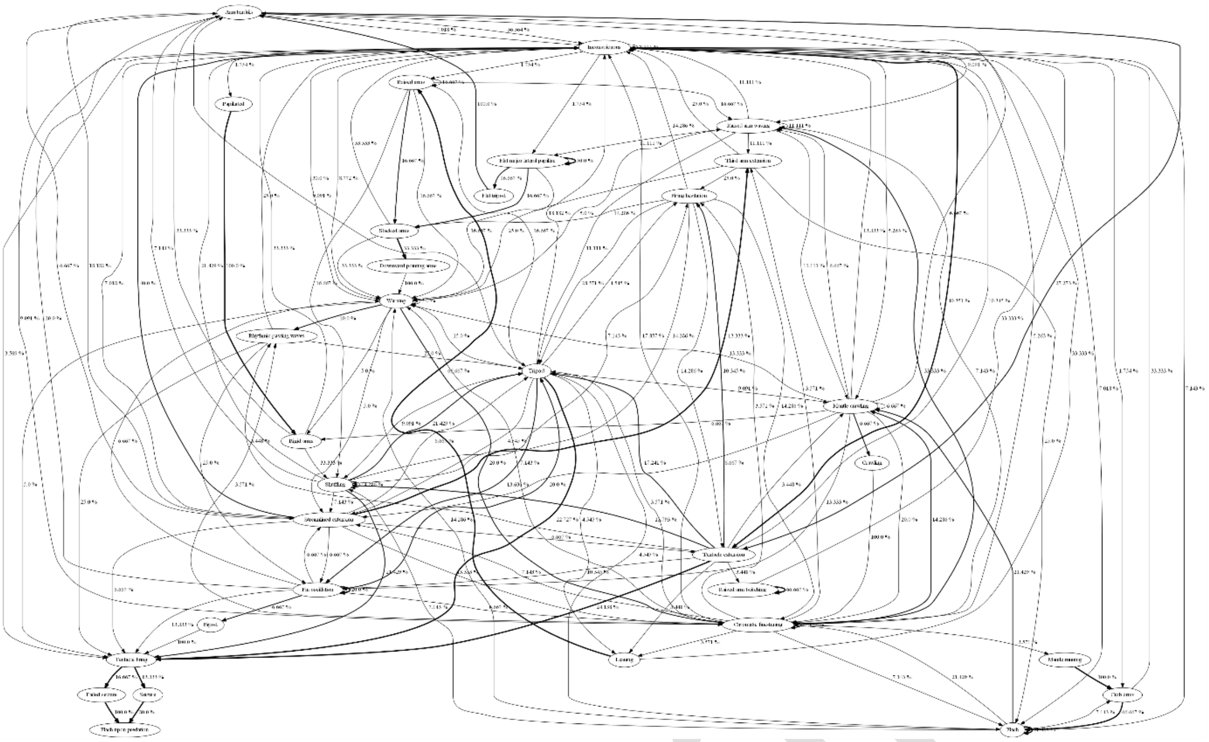

Supplementary Figure S4. All behavioural transitions from all *Sepia apama* observations. Significant behavioural transitions are shown by line thickness ($p = 0.05$ (thick lines) – $p < 0.001$ (thicker lines)).

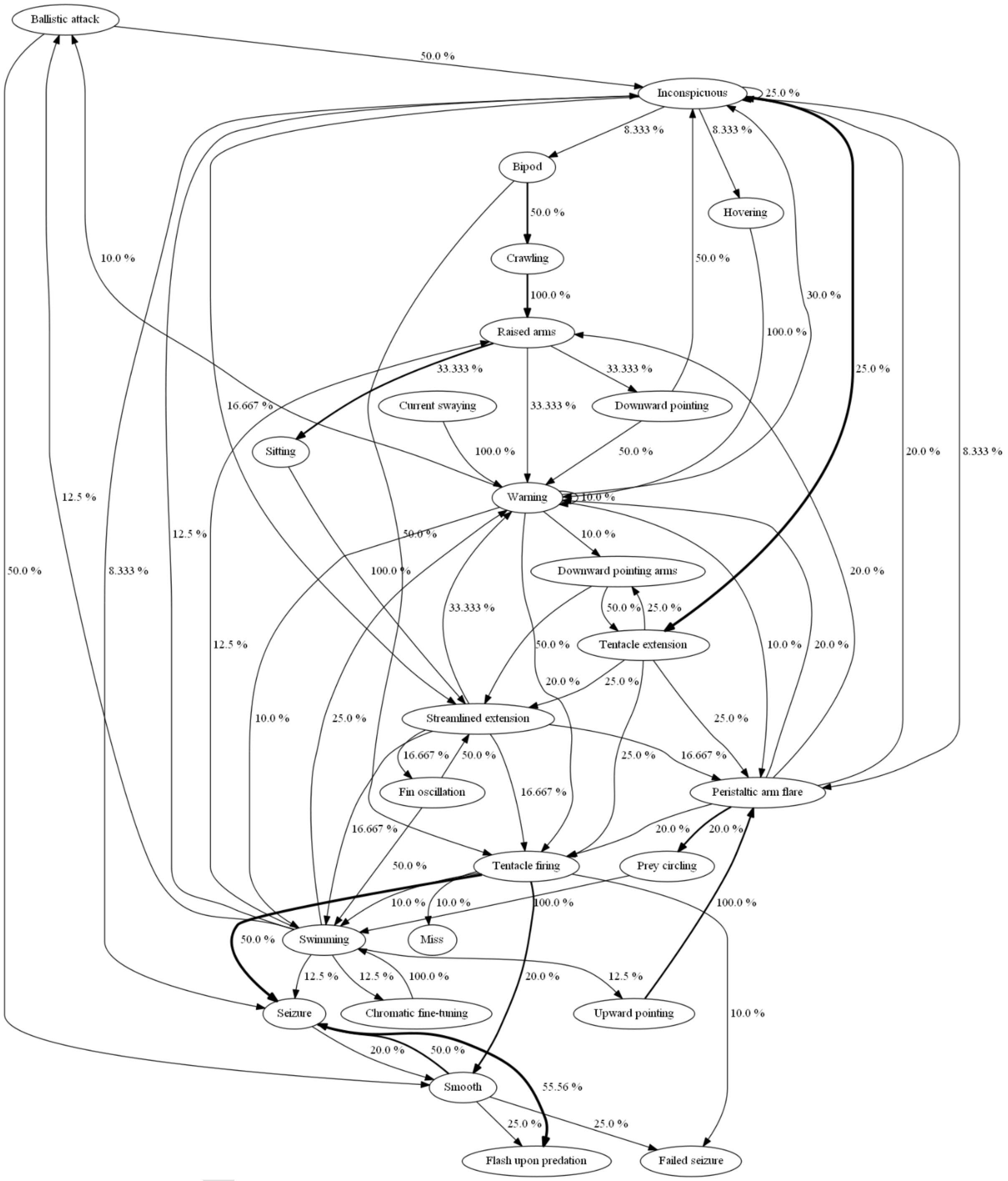


Supplementary Figure S5. All behavioural transitions from all *Sepia latimanus* observations. Significant behavioural transitions are shown by line thickness ($p = 0.05$ (thick lines) – $p < 0.001$ (thicker lines)).

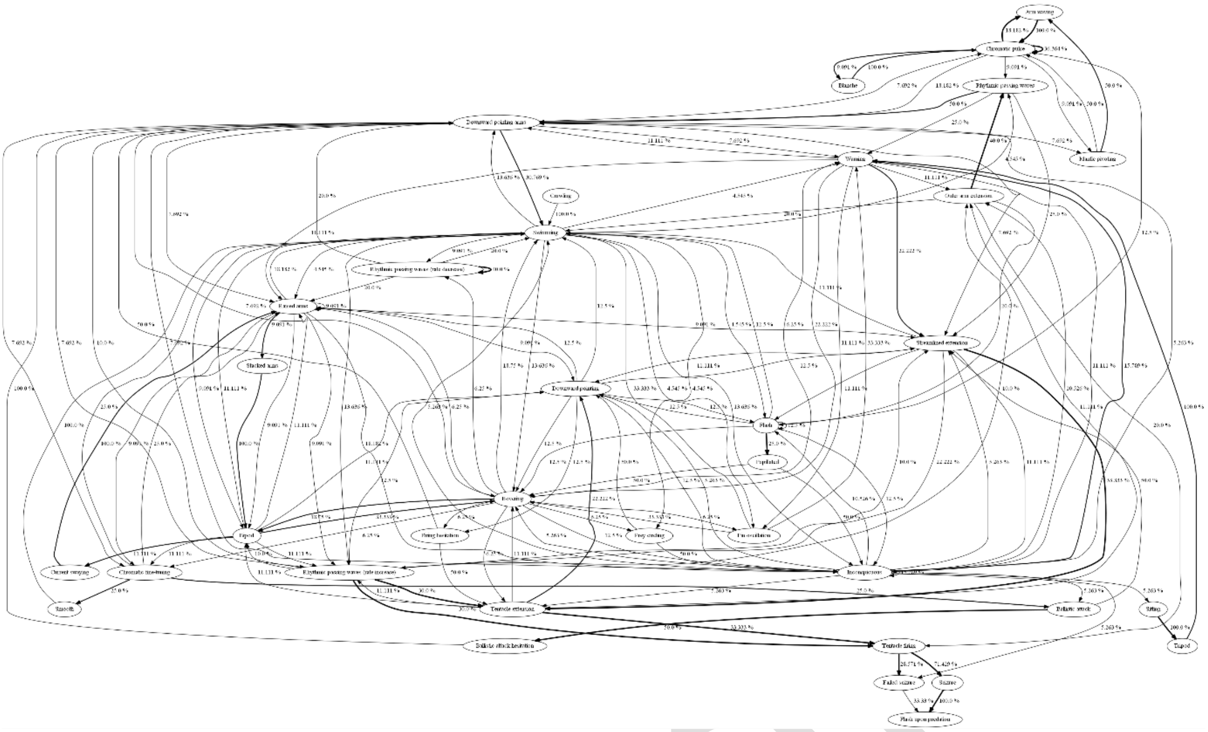

Supplementary Figure S6. All behavioural transitions from all *Sepia officinalis* observations. Significant behavioural transitions are shown by line thickness (p = 0.05 (thick lines) – p < 0.001 (thicker lines)).

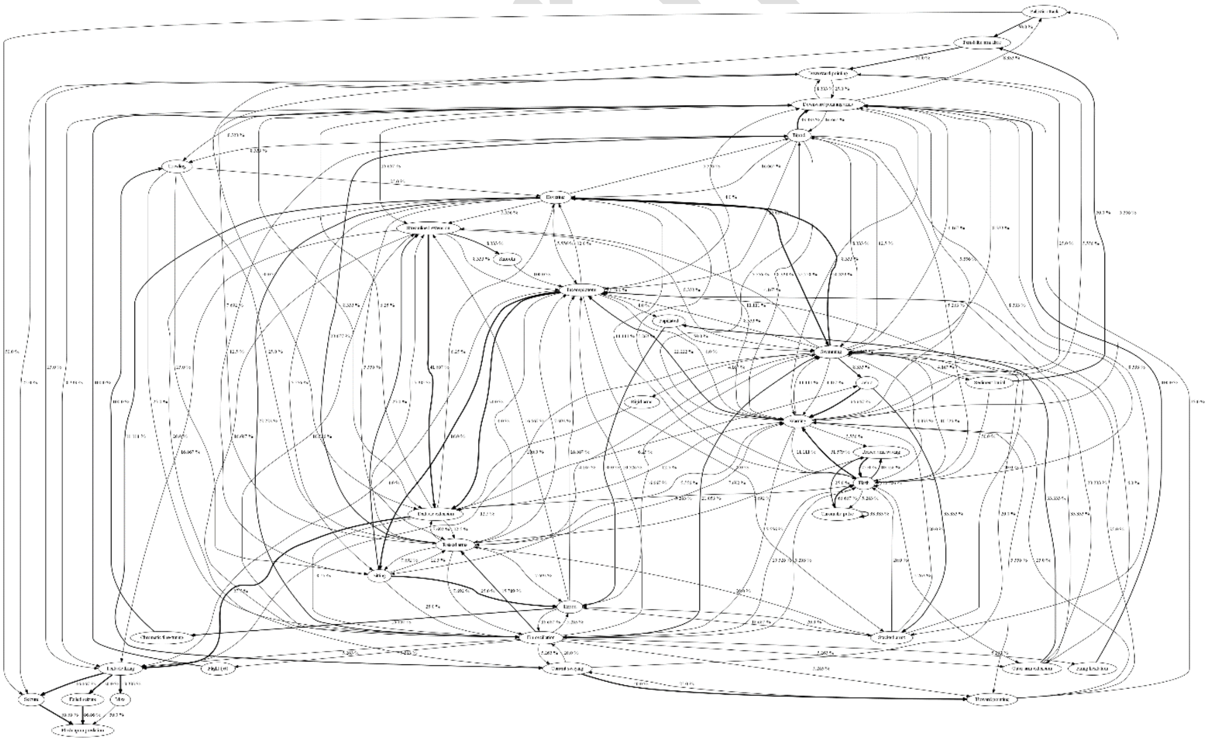

Supplementary Figure S7. All behavioural transitions from all *Sepia pharaonis* observations. Significant behavioural transitions are shown by line thickness (p = 0.05 (thick lines) – p < 0.001 (thicker lines)).

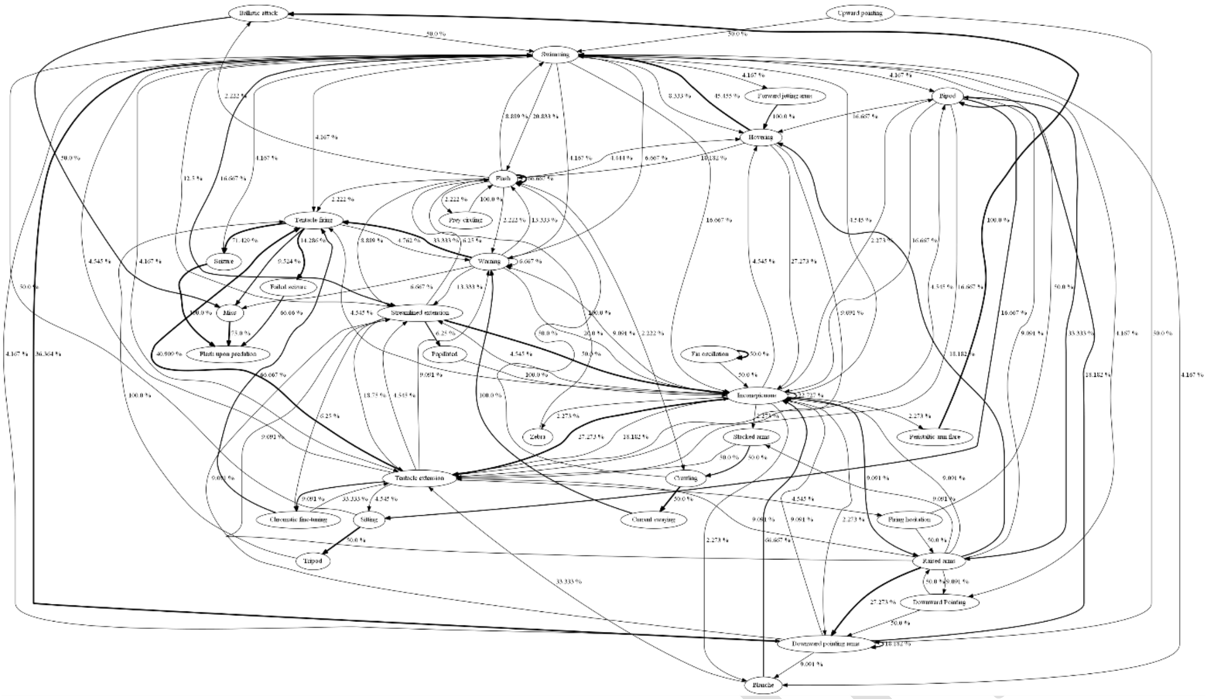

Using social media and citizen science to drive behavioural ecology research: Are cuttlefish using pursuit deterrent signals during hunting?

Dražen Gordon, Philip Pugh and *Gavan M Cooke

Department of Life Sciences, Anglia Ruskin University, Cambridge, United Kingdom.

*Corresponding author

Supplementary Materials "URL list"

Supplementary Table S8. Observation codes (i.e. *M. pfefferi* = Msp, *S. apama* = Sa, *S. latimanus* = Sl, *S. officinalis* = So, *S. pharaonis* = Sp, unspecified *Sepia* species = Sx, various octopus = Oc, and other Decapodiformes = De) for videos analysed in this study

| Observation | URL |
|---|---|
| **Msp1** | https://www.youtube.com/watch?v=UFnooNDn6Z4 |
| **Msp2** | https://www.youtube.com/watch?v=iZNI9dQT-1Y&t=1s |
| **Msp3** | https://www.youtube.com/watch?v=G56x08HCLSw |
| **Msp4** | https://www.youtube.com/watch?v=w1sGxSUnVJI |
| **Msp5** | https://www.youtube.com/watch?v=X0yLJkJqFTY |
| **Msp6** | https://www.youtube.com/watch?v=dYqtfJpIz00 |
| **Msp7** | https://www.youtube.com/watch?v=ab2LLP_rZRk |
| **Msp8** | https://www.youtube.com/watch?v=jTPhY3dykvo |
| **Msp9** | https://www.youtube.com/watch?v=f8OVSfUgCII |
| **Msp10** | https://www.youtube.com/watch?v=--iDZAJbeR0 |
| **Msp11** | https://www.shutterstock.com/video/clip-5924528-flamboyant-cuttlefish---metasepia-pfefferi-hunting |
| **Msp12** | https://www.shutterstock.com/video/clip-5924552-flamboyant-cuttlefish---metasepia-pfefferi-hunting |

| | |
|---|---|
| **Msp13** | https://www.shutterstock.com/video/clip-5924555-flamboyant-cuttlefish---metasepia-pfefferi-hunting |
| **Msp14** | https://www.shutterstock.com/video/clip-15600202-flamboyant-cuttlefish-metasepia-pfefferi-feeding-on-fish |
| **Msp15** | https://www.shutterstock.com/video/clip-17655655-flamboyant-cuttlefish-feeding-metasepia-pfefferi-known-species |
| **Msp16** | https://www.shutterstock.com/video/clip-17655679-flamboyant-cuttlefish-feeding-metasepia-pfefferi-known-species |
| **Msp17** | https://www.shutterstock.com/video/clip-17655706-flamboyant-cuttlefish-feeding-metasepia-pfefferi-known-species |
| **Msp18** | https://www.shutterstock.com/video/clip-18623279-flamboyant-cuttlefish-feeding-on-muck-metasepia-pfefferi |
| **Msp19** | https://www.shutterstock.com/video/clip-18624776-flamboyant-cuttlefish-feeding-on-muck-metasepia-pfefferi |
| **Msp20** | https://www.youtube.com/watch?v=BXb9ZA9XZ58 |
| **Msp21** | https://www.shutterstock.com/video/clip-1011077951-flamboyant-cuttlefish-metasepia-pfefferi-feeding-night-- |
| **Msp22** | https://www.naturefootage.com/video-clips/CW08_021/flamboyant-cuttlefish-feeding |
| **Msp23** | https://www.youtube.com/watch?v=lZVIlldTOkc |
| **Msp24** | https://www.youtube.com/watch?v=--iDZAJbeR0 |
| **Sa1** | https://www.youtube.com/watch?v=MUCduZyCHes |
| **Sa2** | https://www.youtube.com/watch?v=MUCduZyCHes |
| **Sa3** | https://www.youtube.com/watch?v=MUCduZyCHes |
| **Sa4** | https://www.youtube.com/watch?v=MUCduZyCHes |
| **Sa5** | https://www.youtube.com/watch?v=MUCduZyCHes |
| **Sa6** | https://www.youtube.com/watch?v=MUCduZyCHes |
| **Sa7** | https://www.youtube.com/watch?v=MUCduZyCHes |
| **Sa8** | https://www.youtube.com/watch?v=MUCduZyCHes |
| **Sa9** | https://www.youtube.com/watch?v=MUCduZyCHes |
| **Sa10** | https://www.youtube.com/watch?v=MUCduZyCHes |
| **Sa11** | https://www.youtube.com/watch?v=MUCduZyCHes |
| **Sa12** | https://www.youtube.com/watch?v=MUCduZyCHes |
| **Sa13** | https://www.youtube.com/watch?v=AjWX1rGH2SQ |
| **Sl1** | https://www.youtube.com/watch?v=FsqI9p0Z8VU |
| **Sl2** | https://www.youtube.com/watch?v=qjloPutUCeo |
| **Sl3** | https://www.youtube.com/watch?v=jAch1quTkIU |
| **Sl4** | https://www.youtube.com/watch?v=jAch1quTkIU |
| **Sl5** | https://www.youtube.com/watch?v=NuVfLMrpp1g |
| **Sl6** | https://www.naturefootage.com/stock-video-footage?fs=LF04_067 |
| **Sl7** | http://www.footage.net/Search |
| **Sl8** | https://www.youtube.com/watch?v=DXxMWAfLvQA |
| **So1** | https://www.youtube.com/watch?v=nIcFbkpwRmw |
| **So2** | http://www.sarkive.com/invertebrates-marine/sepia-officinalis/video-08a.html |
| **So4** | https://www.youtube.com/watch?v=MjlQ3oQ0rjY |
| **So5** | https://vimeo.com/187549862 |
| **So7** | https://www.naturepl.com/stock-video/common-cuttlefish-(sepia-officinalis)-hunting-filmed-at-night-sark-british/search/detail-0_01474848.html |

| | |
|---|---|
| **So8** | https://www.naturepl.com/stock-video/common-cuttlefish-(sepia-officinalis)-hunting-filmed-at-night-sark-british/search/detail-0_01474842.html |
| **So9** | https://www.youtube.com/watch?v=4qXaPxsa_Wc |
| **So10** | https://www.shutterstock.com/video/clip-43234-sepia-officinalis-hunting-technique-on-plankton-shot |
| **So11** | https://www.shutterstock.com/video/clip-43435-sepia-officinalis-nighttime-shot-captured-wild-mediterranean |
| **So12** | https://www.shutterstock.com/es/video/clip-158704-soon-prey-within-reach-cuttlefish-extracts-retracts |
| **So13** | https://www.shutterstock.com/es/video/clip-6072035-cuttlefish-capture-little-fish-shooting-out-two |
| **So14** | https://www.shutterstock.com/es/video/clip-22780297-cuttlefish-tries-capture-little-fish-shooting-out |
| **So15** | https://vimeo.com/258161890 |
| **So16** | https://www.youtube.com/watch?v=Oe3rlm9UvIg |
| **So17** | https://www.youtube.com/watch?v=xe81UkjxdAE |
| **Sp2** | https://www.youtube.com/watch?v=ykWJBcefp1M |
| **Sp3** | https://www.youtube.com/watch?v=KiUCqOVKEVE |
| **Sp4** | https://www.youtube.com/watch?v=KiUCqOVKEVE |
| **Sp5** | https://www.youtube.com/watch?v=KiUCqOVKEVE |
| **Sp6** | https://www.youtube.com/watch?v=KiUCqOVKEVE |
| **Sp7** | https://www.youtube.com/watch?v=KiUCqOVKEVE |
| **Sp8** | https://www.youtube.com/watch?v=KiUCqOVKEVE |
| **Sp9** | https://www.youtube.com/watch?v=KiUCqOVKEVE |
| **Sp10** | https://www.youtube.com/watch?v=KiUCqOVKEVE |
| **Sp11** | https://www.youtube.com/watch?v=1ptCd1MbP3s |
| **Sp12** | https://www.youtube.com/watch?v=1ptCd1MbP3s |
| **Sp13** | https://www.shutterstock.com/video/clip-1766546-small-cuttlefish-hunts |
| **Sp16** | https://www.youtube.com/watch?time_continue=1&v=Y0oAUkq2zbY |
| **Sp17** | https://www.youtube.com/watch?time_continue=1&v=Y0oAUkq2zbY |
| **Sp18** | https://www.youtube.com/watch?time_continue=1&v=Y0oAUkq2zbY |
| **Sp20** | https://www.naturefootage.com/video-clips/NH05_135/pharaoh-cuttlefish-sepia-pharaonis-feeds-with-feeding-tentacle |
| **Sp21** | https://www.naturefootage.com/video-clips/NH05_135/pharaoh-cuttlefish-sepia-pharaonis-feeds-with-feeding-tentacle |
| **Sp22** | https://www.naturefootage.com/video-clips/NH05_135/pharaoh-cuttlefish-sepia-pharaonis-feeds-with-feeding-tentacle |
| **Sp23** | https://www.naturefootage.com/video-clips/NH05_136/pharaoh-cuttlefish-sepia-pharaonis-feeds-with-feeding-tentacle |
| **Oc1** | https://www.youtube.com/watch?v=Y0o4Wuf1Nt4 |
| **Oc2** | https://www.youtube.com/watch?v=D2oc6HQ3rHQ |
| **Oc3** | https://www.youtube.com/watch?v=jOwRYeLVorM |
| **Oc4** | https://www.youtube.com/watch?v=0nlm-Pp6g64 |
| **Oc5** | https://www.youtube.com/watch?v=3Vm1e9d2uuo |
| **Oc6** | https://www.newsflare.com/video/229538/animals/an-octopus-hunting-a-stone-fish-front-side |
| **Oc7** | https://www2.padi.com/blog/2013/04/03/video-of-the-week-octopus-hunting-prey/ |
| **Oc8** | https://www.huffingtonpost.co.uk/entry/watch-this-tiny-octopus-prove-scientists-wrong_n_55cb5fbbe4b0c3c12bec784 |

| | |
|---|---|
| **Oc9** | https://marinebio.org/two-spot-octopus-hunting-at-night/ |
| **Oc10** | https://www.facebook.com/watch/?v=1020788168028258 |
| **Oc11** | https://www.youtube.com/watch?v=O43hgHGn4KQ |
| **De1** | https://www.youtube.com/watch?v=X2aa6aHEyUI |
| **De2** | https://www.youtube.com/watch?v=5clYS1Z8eQ4 |
| **De3** | https://www.youtube.com/watch?v=okBpSCqrNFA |
| **De4** | https://www.youtube.com/watch?v=Ku4NxKTgShM |
| **De5** | https://www.youtube.com/watch?v=Su6ZCQCzAm4 |
| **De6** | https://www.youtube.com/watch?v=g6XvnCkMpFk |
| **De7** | https://www.express.co.uk/videos/6021578148001/Humboldt-squid-found-hunting-for-fish-at-bottom-of-the-ocean |
| **De8** | https://www.youtube.com/watch?v=AQKs1-fwTgU |
| **De9** | https://vimeo.com/118307814 |
| **De10** | https://www.youtube.com/watch?v=s8XcvqCppTI |
| **De11** | https://www.youtube.com/watch?v=hpgPneFdD4c |
| **De12** | https://www.youtube.com/watch?v=a867k2TkBnE |
| **De13** | https://www.instagram.com/p/B0T4JX7hMew/ |
| **Sx1** | https://www.facebook.com/cephlove/videos/480436916084255/ |

Supplementary Table S9. "Flash upon predation" presence/absence in other cephalopod species.

| Taxa | Prey seizure method | "flash upon predation" | Wild or captive | No. observations |
|---|---|---|---|---|
| **Octopodiformes** | | | | |
| *Hapalochlaena sp.* | Pounce | Yes | Wild | 1 |
| *Octopus cyanea + Plectropomus leopardus* | Speculative hunting | Yes | Wild | 1 |
| *Octopus bimaculoides* | Arm grab | No | Wild | 1 |
| Other sp. | Ballistic attack/ Speculative hunting/Arm grab | Yes | Wild/captive | 8 |
| Other sp. | Ballistic attack/Arm grab | No | Wild | 2 |
| **Decapodiformes** | | | | |
| *Dosidicus gigas* | Tentacle firing | Yes | Wild | 6 |
| *Euprymna scolopes* | Tentacle firing | Yes | Captive | 1 |
| *Sepioloidea lineolata* | Tentacle firing | Yes | Captive | 1 |
| *Idiosepius notoides* | Ballistic attack | Yes | Wild | 1 |
| *Sepioteuthis lessoniana* | Tentacle firing | Yes | Captive | 1 |

Pre-print

| | | | | |
|---|---|---|---|---|
| Shallow water/calamari | Tentacle firing | Yes | Wild | 1 |
| *Sepiola parva* | Tentacle firing | No | Captive | 21 |
| *Sepia mestus* | Tentacle firing | Yes | Wild | 1 |
| *Sepia sp.* | Tentacle firing | Yes | Captive | 1 |

Using social media and citizen science to drive behavioural ecology research: Are cuttlefish using pursuit deterrent signals during hunting?

Dražen Gordon, Philip Pugh and *Gavan M Cooke

Department of Life Sciences, Anglia Ruskin University, Cambridge, United Kingdom.

*Corresponding author

Supplementary materials “unusual behaviours in cuttlefish”

Supplementary Table S10. Unusual cuttlefish behaviours witnessed from social media and citizen science videos analysed.

| **Species** | **Category** | **Description** | **Interpretation/context** |
|---|---|---|---|
| | Cephalopod predation | | |
| *S. plangon* | Cephalopod predation | *S. plangon* preyed upon *Sepioloidea lineolata* | Nutrition |
| | https://www.instagram.com/p/Bz6ac3UhkhN/ | | |
| | Hunting strategy | | |
| *M. pfefferi* | Social hunting | Male and female pair forage and hunt different prey together | Mating pair, or more efficient prey-spotting |
| | https://www.youtube.com/watch?v=8DtBp7LU07Q | | |
| *S. latimanus* | Clouding prey | Oscillating fins and moving arms disturb substrate. Disturbed substrate hits benthic prey | Attempt to confuse prey, or reduce prey vision |
| | https://www.youtube.com/watch?v=emWk9Or0GVU | | |
| | Behaviours | | |
| *M. pfefferi* & *Sepia spp.* | Pigmented tentacular clubs | Feeding tentacles are not uniformly pale, clubs are coloured | Directive mark, draws prey attention |
| | https://www.naturefootage.com/video-clips/CW08_021/flamboyant-cuttlefish-feeding | | |
| *M. pfefferi* & *Sepia spp.* | Flash upon predation | Abrupt brightening or change in body pattern. Can include whole body or body parts | Pursuit-deterrent signal, dissuades potential threat |
| | https://www.shutterstock.com/es/video/clip-6072035-cuttlefish-capture-little-fish-shooting-out-two | | |

| | | | |
|---|---|---|---|
| *M. pfefferi* | Arm tendrils | Mechanically wriggled tendril-like projections extending from third arm pair | Directive mark, draws prey attention |
| *M. pfefferi* | Flat tripod<br>https://www.youtube.com/watch?v=X0yLJkJqFTY | Fourth arm pair are completely extended, head and ventral portion of mantle are in complete contact with substrate | Enables substrate level tentacular firing |
| *M. pfefferi* | Crab arms | Arms possibly mimic crustacean claws and antenna | Attempt to deceive predators/prey |
| *M. pfefferi* | Crab head | Pigmentation resemblant of a crustacean (orange, black, yellow) – expressed with "crab arms" | Attempt to deceive predators/prey |
| *M. pfefferi* | Animation eyes<br>https://www.youtube.com/watch?v=--iDZAJbeR0 | White eye patches with thick black outlines | Conceals eyes |

### Using social media and citizen science to drive behavioural ecology research: Are cuttlefish using pursuit deterrent signals during hunting?

Dražen Gordon, Philip Pugh and *Gavan M Cooke

Department of Life Sciences, Anglia Ruskin University, Cambridge, United Kingdom.

*Corresponding author

Supplementary materials "Supplementary discussion"

We found that retroactively gathered citizen science and unsolicited social media observational data provide a vital tool for studying the behavioural ecology of hunting in five Sepiidae species.

*S. apama* increases streamlining during ballistic attack events by shifting from "papillated" to "smooth" epithelial texture. Arm postures (e.g. *S. apama*, *S. latimanus*, *S. officinalis*) or body patterns (e.g. *S. apama*, and *S. latimanus*) were also adapted during ballistic attacks (see Supplementary Figure S4, S5, & S6).

Warning displays signalled by species studied here included, 'deimatic displays' (and constitutive components i.e. 'paired mantle spots', 'mantle margin stripe', etc)[1] (see Supplementary Table S1) and 'intense zebra patterning' (e.g. *S. officinalis* and *S. pharaonis*) (see Supplementary Table S7, So2 & So15) – zebra patterning is usually considered to be exclusively used towards competing male congeners during courtship events[2–4]. Cuttlefish species also performed dynamic patterns: 'flash' (all species studied here except *S. apama*), 'chromatic pulse' (e.g. *S. latimanus* and *S. officinalis*), and 'rhythmic passing waves' (e.g. *M.*

Pre-print

*pfefferi* and *S. latimanus*); which have been speculated to have aposematic purposes in cuttlefish[5], but also in other taxa (i.e. fish, crustacea, and polychaetes use bioluminescent flashing as an aposematic signal)[6].

Aposematic (*M. pfefferi*) or deimatic warning displays (*Sepia* spp.) seen in cuttlefish here (Figure 3) may not always deter predation from all predator species, and Langridge[7] have provided evidence that *S. officinalis* selectively avoids signalling deimatic displays towards larger, more dangerous predators, like the smooth hound (*Mustelus mustelus*). This may explain why despite being thought to be poisonous, and using conspicuous aposematic warning displays, *M. pfefferi* also employs a flash upon predation signal.

When searching for useful videos on social media we observed "flash upon predation" to be used upon prey seizure by other coleoid taxa as well. Examples include: blue-ringed octopus (*Hapalochlaena sp*.) (see Supplementary Table S8 & S9) and the social hunting (see Benoit-Bird & Gilly[8]; Vail & Bshary[9] for evidence of social hunting per se) day octopus (*O. cyanea*) - *O. cyanea* hunts cooperatively with coral trout (*Plectropomus leopardus*); (see Benoit-Bird & Gilly[8]) and Humboldt squid (*Dosidicus gigas*) (see Supplementary Table S8 & S9).

1. Hanlon, R. T. & Messenger, J. B. Cephalopod Behaviour Cambridge University Press. *Cambridge, UK* (1996).

Pre-print
